# Cryptic Cryptophytes – revision of the genus *Goniomonas*

**DOI:** 10.1101/2024.07.17.603845

**Authors:** Maria Sachs, Frank Nitsche, Hartmut Arndt

## Abstract

Cryptomonad protists are ubiquitously distributed over marine and freshwater habitats. As an exception to the coloured cryptomonads, the heterotrophic cryptomonads of the genus *Goniomonas* have an ancestral phylogenetic position. They lack any kind of chloroplast and most likely represent a basal group to those cryptomonad groups having obtained their chloroplast by secondary endosymbiosis. Earlier studies have shown a deep divergence between freshwater and marine clades of goniomonads that comprise large genetic distances between members within the group and also between the two groups of marine and freshwater taxa. Still, marine and freshwater species carry the same genus name, and until up to date only few species have been described. We therefore restructured goniomonad systematics based not only on a separation of marine and freshwater taxa, but, taking the large genetic distances into account, also several new genera are described. Based on morphological as well as phylogenetic data (18S rDNA sequences), this leads to the formation of the freshwater genera *Limnogoniomonas* n. g., *Goniomonas* and *Aquagoniomonas* n. g. and the marine genera *Neptunogoniomonas* n. g., *Baltigoniomonas* n. g., *Marigoniomonas* n. g., *Thalassogoniomonas* n. g., *Poseidogoniomonas* n. g. and *Cosmogoniomonas* n. g. To give the restructuring process a stable basis, we additionally propose a neotype for *Goniomonas truncata*.

## INTRODUCTION

Cryptomonad flagellates are small, mostly bean shaped, non-colonial protists that encompass at least 200 species (Novarino, 2003). The group is globally distributed in marine, brackish and freshwater habitats (Archibald, 2020). Cryptomonads played a crucial role in understanding evolutionary processes as phototrophic representatives are the outcome of secondary endosymbiosis (Cenci et al., 2018; McFadden, 2018). Phototrophic cryptomonads had engulfed red algae which themselves were originally created through the engulfment of a cyanobacterium by a eukaryotic phagotroph during primary endosymbiosis (Keeling, 2004). To understand the evolutionary history of cryptophytes and elucidate their unique characteristics and adaptations, Douglas and Penny (1999) provided important insights into the evolutionary dynamics of their plastid genomes by emphasizing the descent of their plastids from red algae and the retention of nucleomorphs as a distinguishing feature among secondary plastid-bearing algae (Douglas and Penny, 1999). Within the larger group of cryptists, which include *Palpitomonas bilix* and kathablepharids (as evolutionary more basal groups), as well as cryptophytes and goniomonads, it is a likely scenario that photosynthesis was established after the split of the latter two (Yabuki et al., 2014). This would potentially make goniomonads a last form of pre-endosymbionts before the engulfment of plastids. Cenci et al. (2018) sequenced the entire genome of the most recently described goniomonad species *Goniomonas avonlea*. On genomic or phylogenetic level, no convincing evidence could be found that *G. avonlea* ever possessed a plastid supporting the theory that *Goniomonas* is indeed a very basal phagotroph, a “pre-secondary-endosymbiosis cryptomonad” (McFadden, 2018). Apart from the lack of plastids, all goniomonads share a number of morphological traits, such as a laterally flattened, anteriorly more or less truncated cell body and a horizontal row of ejectosomes in the anterior part of the cell (Novarino, 2003). Unlike other cryptomonads which were reported to have a “recoiling” swimming behaviour (Novarino, 2003), goniomonads display a rather “jerky” movement (Kim and Archibald, 2013), and move on the substrate along irregular tracks while rotating. When analysing sediment associated protists, *Goniomonas*-like flagellates are among the 20 most commonly observed heterotrophic flagellate morphotypes (Patterson and Lee, 2000) and have been recorded in many different habitats (Arndt et al., 2000). Until the taxon was transferred to the currently used named genus *Goniomonas* by Stein (1878), it was considered under the protonym *Monas truncata* (Fresenius, 1858). Larsen and Patterson (1990) discussed the history of using the genus name in the past and described two new marine species of *Goniomonas*. Up to now, five different species have been described, with *G. avonlea* (Kim and Archibald, 2013) being the most recent. Apart from the type species *G. truncata* and *G. brasiliensis* (Jarreta de Castro and Bicudo, 2023) which were isolated from freshwater, the three other species are marine species, *G. amphinema*, *G. pacifica* (see Larson and Patterson, 1990) and *G. avonlea* (Kim and Archibald, 2013).

Recently, the parallel study of Phanprasert et al. (2025) added another three marine species, *G*. *ulleungensis, G. lingua* and *G. duplex* to the list of scientifically described species. Molecular phylogenetic analyses revealed a deep divergence between the goniomonad species from marine and freshwaters (von der Heyden et al., 2004). Studies on other protist groups have shown similar differences in ecological clusters associated with molecular differentiations, for example the choanoflagellate taxa *Codosiga* and *Hartaetosiga* (Carr et al., 2017). The high divergence among the genus *Goniomonas* could imply that there are still multiple undescribed species and that the genus is generally under-split from a taxonomical point of view (von der Heyden et al., 2004). The taxonomic situation becomes even more complex since type material other than drafts of microscopic images are missing. Studies by Shalchian-Tabrizi et al. (2008) and Shiratori and Ishida (2016) have already indicated a large hidden diversity within cryptomonads, including heterotrophic lineages.

In this study, we describe seven new species and emend several other strains from different aquatic habitats including marine, brackish and freshwater environments and propose the revision of the genus *Goniomonas* into *Goniomonas* sensu strictu, which includes freshwater species and a newly created neotype for *Goniomonas truncata*, as well as five new genera within both the freshwater and marine clusters based mainly on genetic distances, including also the recently described species of Phanprasert et al. (2025). Morphological descriptions are supported by light and electron micrographs and genetic divergences based on 18S rDNA phylogeny.

## MATERIALS AND METHODS

Sediment and plankton samples were taken at diverse sampling sites (Supplement Table 8). Sediment samples were taken with a Multi-Corer system (MUC), while plankton samples were either taken with a CTD-rosette system from different depth or were taken with flasks from surface waters. Subsamples of a few millilitres were suspended in ambient autoclaved water (approx. 20 ml) and transferred to 50-ml tissue culture flasks (Sarstedt, Nümbrecht, Germany). Isolation of cells from raw cultures was carried out by micromanipulation under an inverted microscope or by the liquid aliquot methods pipetting a few microlitres of raw cultures in either 0.8- or 0.4-ml multi-well plates containing ambient autoclaved water. Cultures flasks and wells were supplied with autoclaved Quinoa grains to support the growth of autochthonous bacteria.

Mono-cultures of strains HFCC (**H**eterotrophic **F**lagellate Culture **C**ollection **C**ologne at the University of Cologne) 22, 157, 272, 841 and 1666 with marine/brackish origin were further cultivated in Schmalz-Pratt medium (Suppl. Table 1). Salinity was adjusted to original salinity of the sampling site. Freshwater strains HFCC 235, 251, 23, 5007, 162 were cultivated in WC medium (Guillard and Lorenzen, 1972). To stimulate growth of associated bacteria, either autoclaved wheat or quinoa grains were added to the cultures as a carbon source.

### Morphological analyses

Light microscopical investigations were carried out using an inverted microscope (Zeiss Axio Observer; water immersion condenser; 100x/1.4NA oil immersion objective and differential interference contrast). High resolution videos were recorded from cultures in Petri dishes with coverslip using two ways. Earlier records were based on an Allen Video-enhanced system (Hamamatsu C6489, Argus-20) to supress noise and amplify contrast. Frame by frame evaluation of videos was carried out using VirtualDub (www.virtualdub.org) and editing by ImageJ (Abramoff et al., 2004). Morphological measurements were carried out using Axio Vision Rel. 4.8 (Zeiss, Germany). Observations of recent isolates were made using videos recorded with the camera Axiocam 305 mono 31812 (Zeiss, Germany). Pictures were analyzed using the ZEISS ZEN 3.5 blue edition program.

For electron microscopical investigations, cultures were fixated with 0.1 M cacodylate buffered glutaralaldehyde solution (2.5%) for one hour at 4°C. Fixated cells were washed twice with cacodylate buffer and once with distilled water and then stained with 0.5% osmium tetroxide solution for ten minutes. After two additional washing steps with distilled water, the cells were step-wise dehydrated with an increasing ethanol series (30, 50, 70, 80, 90, 96%; each step 15 min) followed by ethanol/HMDS (1:1) and dried in pure HDMS (Hexamethyldisilazan, Carl Roth, Germany). Before examination by SEM (Fei Quanta 250 FEG at the Biocenter, University of Cologne), cells were sputtered with a gold layer (12 nm).

### Molecular studies

Molecular analyses followed procedures described by Schoenle et al. (2020). Strains were centrifuged (4000 x g for 20 min at 4°C, Megafuge 2.0 R, Heraeus Instruments) and genomic DNA was extracted using the Quick-gDNA^TM^ MiniPrep (Zymo Research, USA). For sequencing partial SSU rDNA, three different sets of primers were used: 18S-For (5’- AACCTGGTTGATCCTGCCAGT 3’) and 18S-Rev (5’TGATCCTTCTGCAGGTTCACCTAC 3’) (Medlin et al., 1988), 590For (5’-CGGTAATTCCAGCTCCAATAGC-3’) in combination with 1300 Rev (5’-CACCAACTAAGAACGGCCATGC-‘3) (Wylezich et al., 2002) and the set 82F (5’-GAAACTGCGAATGGCTC-3’) (López-García et al., 2002) in combination with Rev-1 (5’-ACCTACGGAAACCTTGTTACG-3’) (newly assigned) were used for amplification with a primer concentration of 1 µM and the help of a PCR Mastermix (VWR Life Science, Red Taq DNA Polymerase, Hassrode, Belgium). For the first two primer sets the following steps were applied: initial denaturation step at 98°C for 2 min, followed by 35 cycles of 98°C for 30 s, 55 °C for 45 s and 72°C for 2.30 min and a final elongation step for 10 min at 72°C. For the third primer set, the following steps were used: initial denaturation at 98°C for 2 min, 35 cycles at 98°C for 30 s, at 52°C for 45 s, at 48°C for 30 s and at 72°C for 2.30 min, final elongation step at 72°C for 10 min. Length of PCR products were checked on a 1% agarose gel and subsequently purified with the PCR Purification Kit (Jena Bioscience, Jena, Germany). Purified PCR products were sequenced at GATC Biotech, Germany.

### Phylogenetic analyses

The 18S rDNA of the 26 new goniomonad sequences were aligned with 25 closely related *Goniomonas* sequences from GenBank, and the tree most recently described strains by Phanprasert et al. (2025), as well as six outgroup sequences of the CRY1 group (following Shiratori and Ishida, 2016) that includes both freshwater and marine strains, as well as *Hemiarma marina*. Sequences were aligned using the MAFFT algorithm (Katoh and Standley, 2013) and were slightly manually corrected with the help of BioEdit (Hall, 1999). The corrected alignment was analysed with the Maximum likelihood (ML) method using RAxML v.8.2.12 (Stamatakis, 2014) applying the (GTR) model + Γ + I model of rate heterogeneity with 1000 replicates. Pairwise distances were calculated with BioEdit (Hall, 1999). Bayesian analysis was carried out with MrBayes on XSEDE (3.2.7a) tool (Huelsenbeck and Ronquist, 2001; Ronquist et al., 2012) using the (GTR) model + Γ + I with a sampling frequency for the Markov chain set to 10. The stop value of topological convergence diagnostic was set to 0.01. The number of generations/cycles for the MCMC algorithm was set initially to 1,000,000 and was reached after 256,000 generations. Unalignable sites were treated as missing data. The final version of the tree was edited using MEGA-X (Kumar et al., 2018), FigTree (Version 1.4.4, https://github.com/rambaut/figtree/releases), and Inkscape (Version 1.0.1) (https://www.inkscape.org/). P-distances were calculated by aligning two sequences and after cutting the unaligned front and back sequence sites, they were aligned again and the individual distance was calculated in percent.

### Synapomorphy analysis

To identify synapomorphic sequence motifs that were shared within the phylogenetic clades, we used a custom Python script built on the Biopython library (Cock et al., 2009) to screen the alignment. The script detected unique, non-overlapping motifs in the conserved regions of the alignment and occurred in less than 10 % of sequences outside the clade. The detected motifs were carefully checked manually to avoid redundancy. Similarities as well as differences were visualized using Inkscape (Version 1.0.1) (https://www.inkscape.org/). Differences in nucleotides within the clades were shown using the IUPAC letter code.

## RESULTS

### Morphology

All “*Goniomonas*-like” strains investigated in this study shared the following morphological characteristics: an anteriorly truncated cell body, rounded at the posterior end, cell covered with periplast plates, two flagella arising laterally at the cell anterior and a band of ejectosomes appearing transversely at the anterior end of the cell. Those basic characteristics varied especially between the freshwater and marine clade, with freshwater strains generally being larger in cell size and more elongated in shape, whereas marine strains were overall smaller and more roundish (Table 1). A contractile vacuole is generally located on the ventral side below the band of ejectosomes in the anterior part of cells, clearly visible in freshwater strains, but less visible in marine strains. For determination of the ventral and dorsal side of the cell, we decided to follow the terminology of Mignot (1965) that designates the narrow side, where the flagella emerge as the dorsal surface and the opposite narrow site as the ventral surface. The two wide sides are determined as left and right sides (Figure 1).

**Figure 1.**
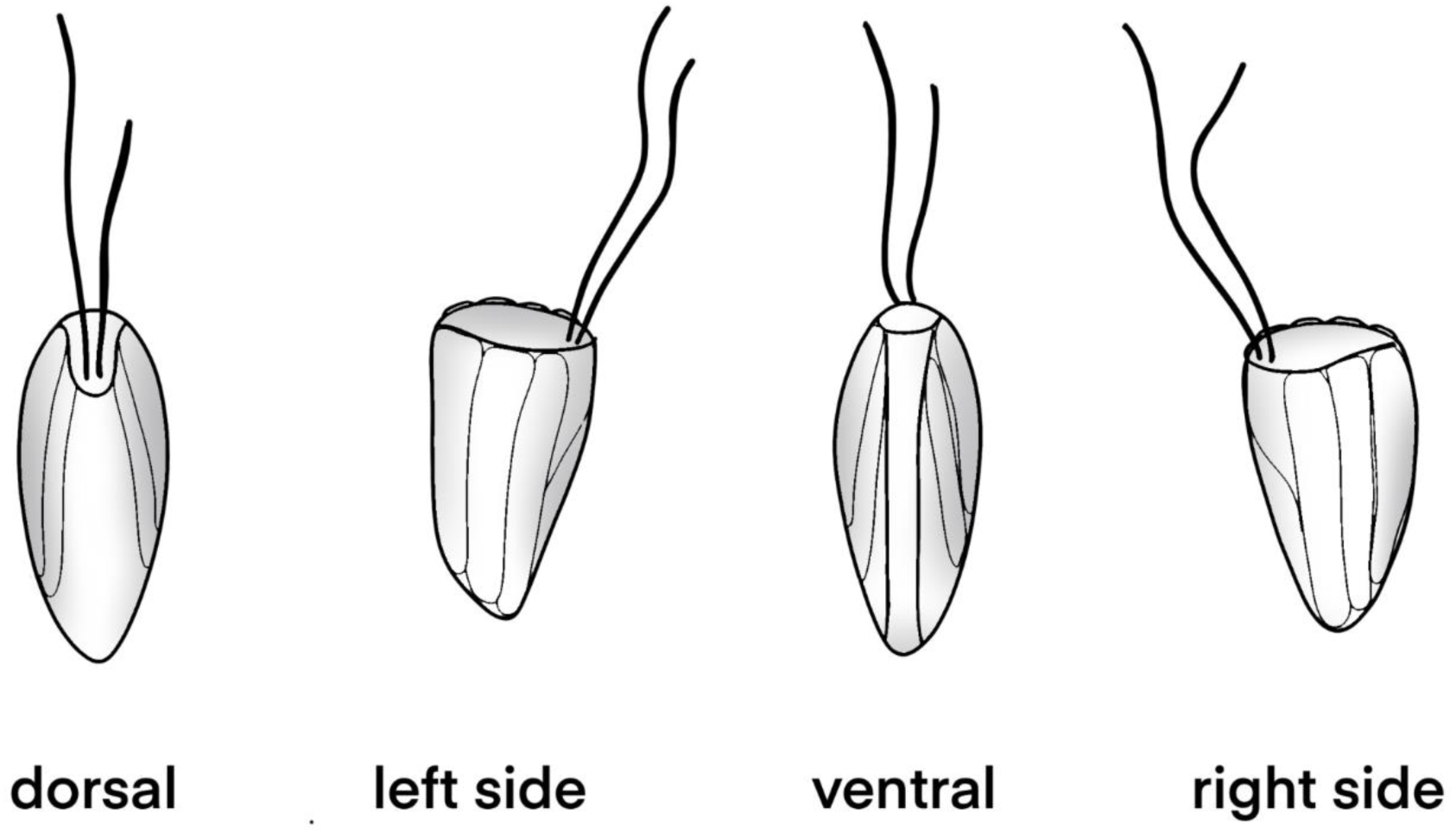
Illustration of cell orientation for goniomonads following the terminology of Mignot (1965).

**Table 1.**
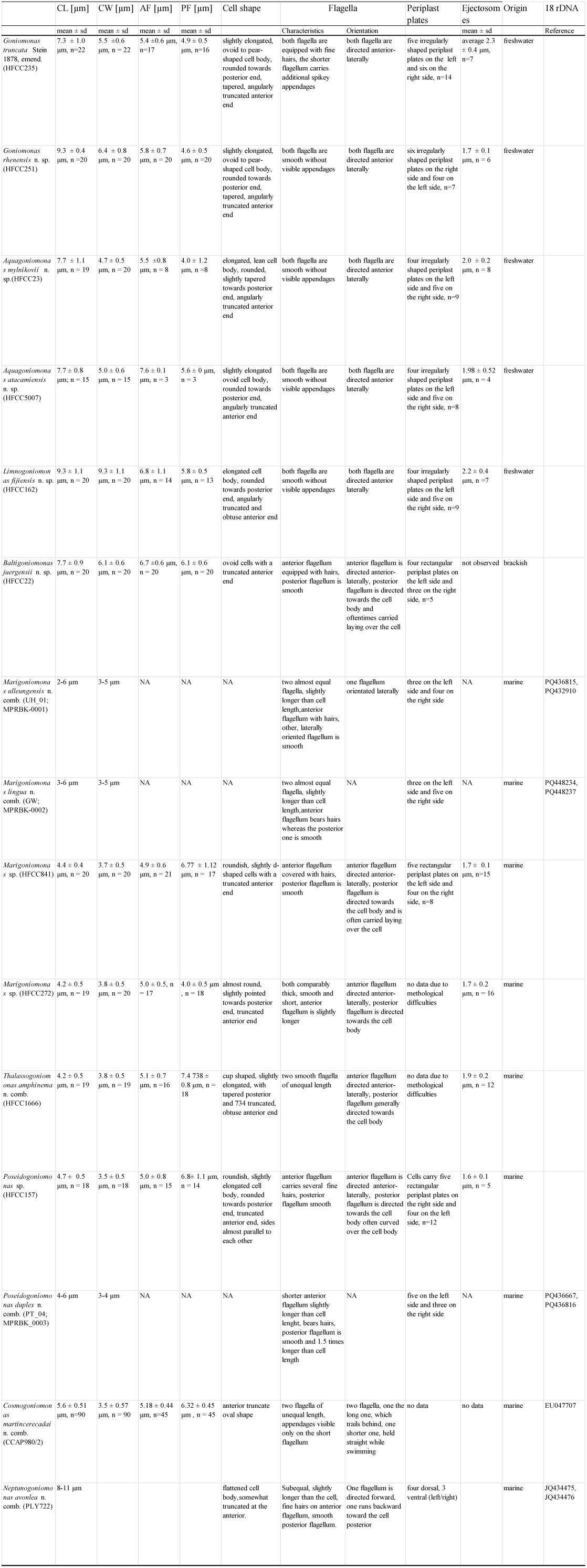
Landscape table of morphological characteristics summarized for new species and strains investigated in this study. CL: cell length, CW: cell width, AF: anterior flagellum, PF: posterior flagellum.

Two out of ten analysed strains for this study (HFCC 235 and 251) could be assigned to the genus *Goniomonas* (Figure 2, A-G resp. H-K) isolated from freshwater. The cell shape was always laterally compressed, slightly elongated ovoid and sometimes pear shaped (HFCC 251). The elongated shape makes the left and right margin of the cells appearing almost parallel Two other goniomonad strains (HFCC 23 and 5007) were assigned to the newly designed freshwater clade *Aquagoniomonas* n. g. (Figure 2, L-P resp. Q-T, Figure 3). Morphologically similar, though genetically distinct were the following clades: Strain HFCC 162 (Figure 4) was assigned to the new genus *Limnogoniomonas* n. g. Morphological observations for strain HFCC 22 assigned to the genus *Baltigoniomonas* n. g. (Figure 5, A-D) show a similar cell size as *Limnogoniomonas* n. g. Both strains are free living and display a similar swimming behaviour like *Goniomonas* and *Aquagoniomonas* n. g. Cells are laterally compressed and slightly elongated. They carry two flagella of unequal length of which one possesses stiff flagellar hairs. Strains HFCC 841 and 272 both fall into the newly created genus *Marigoniomonas* n. g. (Figure 5, E-I and J-M). In relation to the cell size, the flagella are much longer compared to the freshwater strains. Both strains show a roundish rather compact cell shape, not elongated in contrast to the freshwater genera. Cells of strain HFCC 841, which were isolated from the Chilean Pacific coast near Antofagasta, were 3.6–4.9 µm (average 4.4 ± 0.4 µm, n = 20) long and 2.9–4.6 µm (average 3.7 ± 0.5 µm, n = 20) wide. Flagella were of unequal length with the anterior flagellum being approx. 1/3 shorter with 4.2–6.3 µm (average 4.9 ± 0.6 µm, n = 21) length and covered with hairs, while the posterior flagellum was smooth and directed towards the cell body and is often carried laying over the cell, 5.1–8.5 µm (average 6.77 ± 1.12 µm, n =17) long. Cells carry five rectangular periplast plates on the left side and four on the right side. Periplast plates could not be analysed for strain HFCC 272 due to difficulties with the preparation. Cells of strain HFCC 272, which were isolated from the Atlantic coast of Madeira from 950m depth off Ribeira, were 3.3–5.0 µm (average 4.2 ± 0.5 µm, n = 19) long and 3.1–5.0 µm (average 3.8 ± 0.5 µm, n = 20) wide. The anterior flagellum was slightly longer and directed anterior-laterally with 5.0 ± 0.5 µm (4.1–5.7 µm, n = 17) length, the posterior flagellum was directed towards the cell body and was 3.2–4.8 µm (4.0 ± 0.5 µm , n = 18) long. In addition to the prominent band of ejectosomes, a smaller nodule like ejectosomes between the periplast plates was visible.

**Figure 2.**
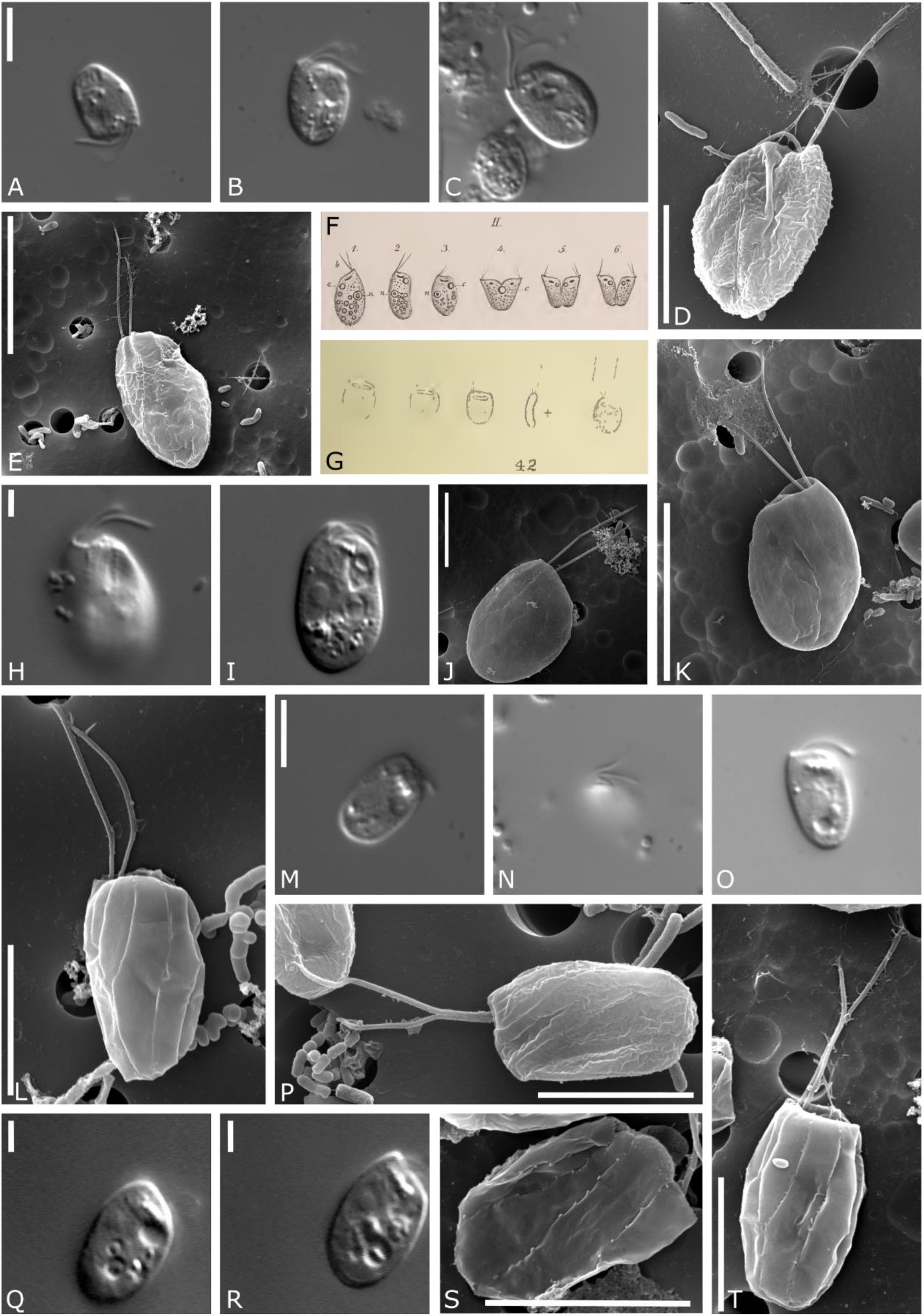
Light and electron micrographs and drawings of the genera *Goniomonas* and *Aquagoniomonas* n. g. et n. sp. A-E *Goniomonas truncata* (bar indicates: A, D-5 µm, E-10 µm), F drawing of *Goniomonas truncata* (from Stein, 1878), G drawing of *Monas truncata* (from Fresenius, 1858), H-K *Goniomonas rhenensis* n. sp. (bar indicates: H-I 2µm, J-5 µm, K-10 µm), L-P *Aquagoniomonas mylnikovii* n. g et. n. sp. (bar indicates: L, M, P-5 µm), Q-T *Aquagoniomonas atacamiensis* n. g. et n. sp. (bar indicates: Q, R-2 µm, S, T-5 µm).

**Figure 3.**
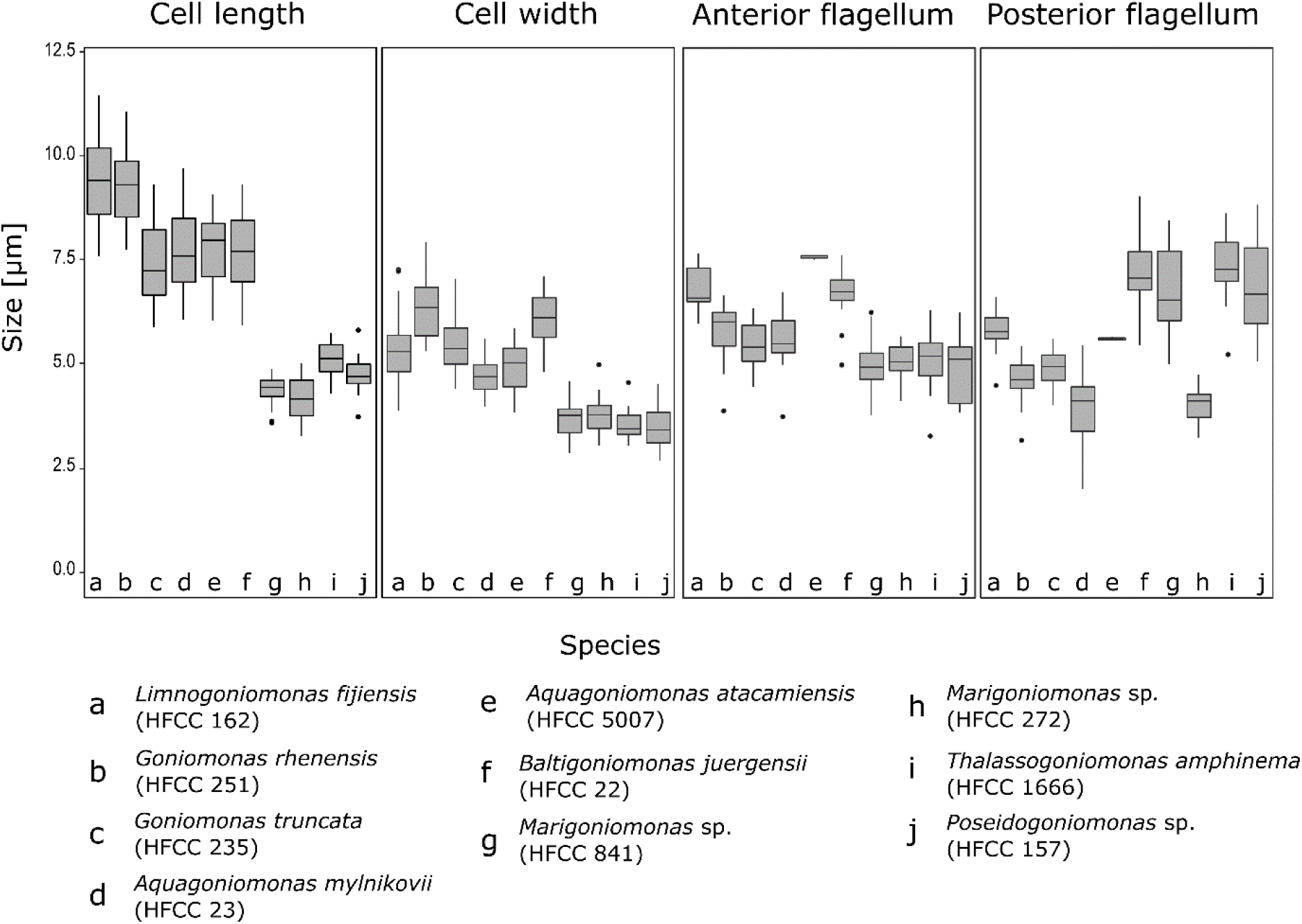
Boxplots showing the comparison of morphological measurements of the described goniomonads. Letters in the box plots indicate the different species. a: *Limnogoniomonas fijiensis* n. g. et n. sp.; b: *Goniomonas rhenensis* n. sp.; c: *Goniomonas truncata*, d: *Aquagoniomonas mylnikovii* n. g. et n. sp., e: *Aquagoniomonas atacamiensis* n. g. et n. sp.; f: *Baltigoniomonas juergensii* n. g. et n. sp.; g: *Marigoniomonas* sp. (HFCC 841); h: *Marigoniomonas* sp. (HFCC 272); i: *Thalassogoniomonas amphinema* n. comb.; j: *Poseidogoniomonas* sp. (HFCC 157)

**Figure 4.**
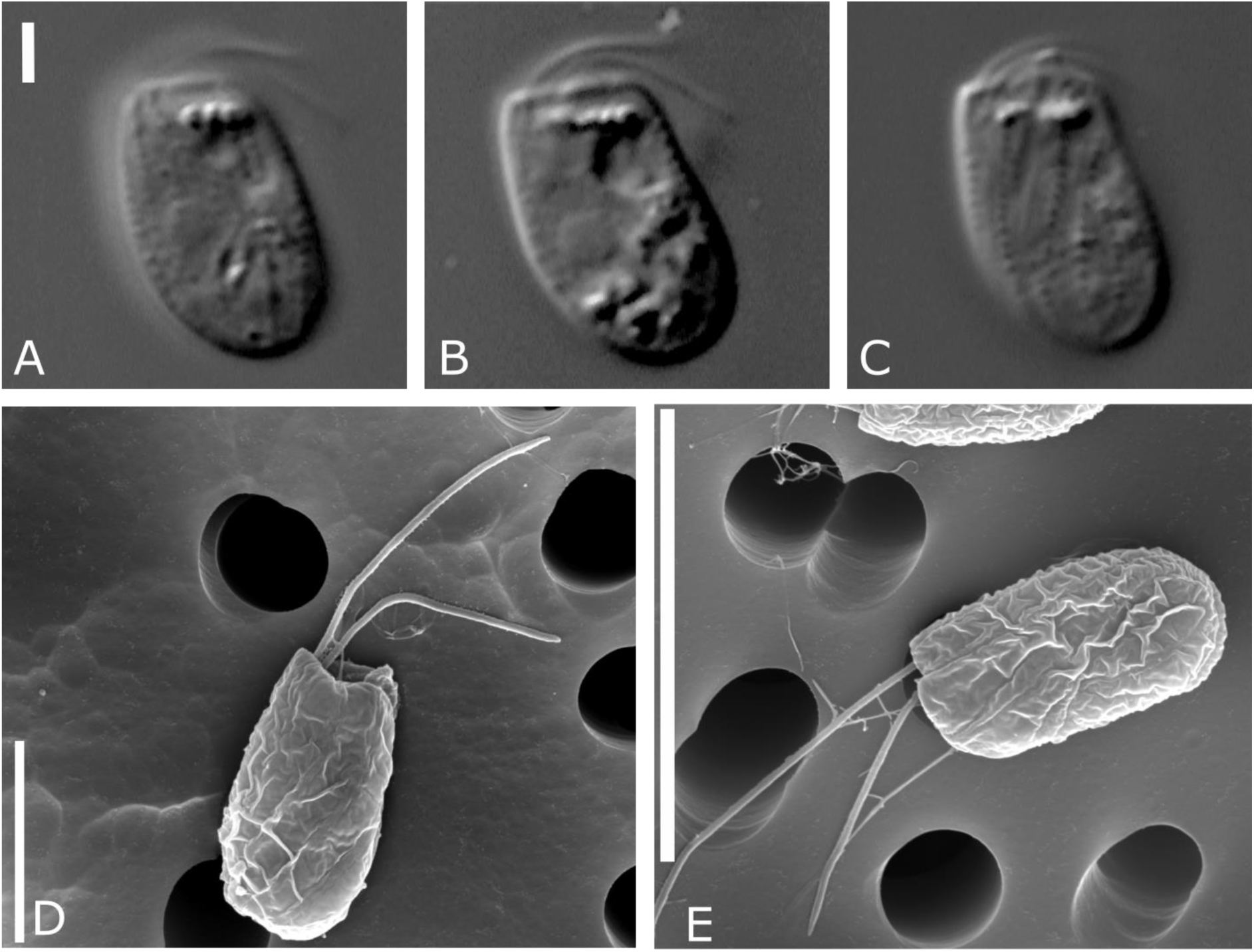
Light and electron micrographs showing *Limnogoniomonas fijiensis* n. g. et. n. sp. (A-E; bar indicates: A-2 µm, D-5 µm, E-10 µm)

**Figure 5.**
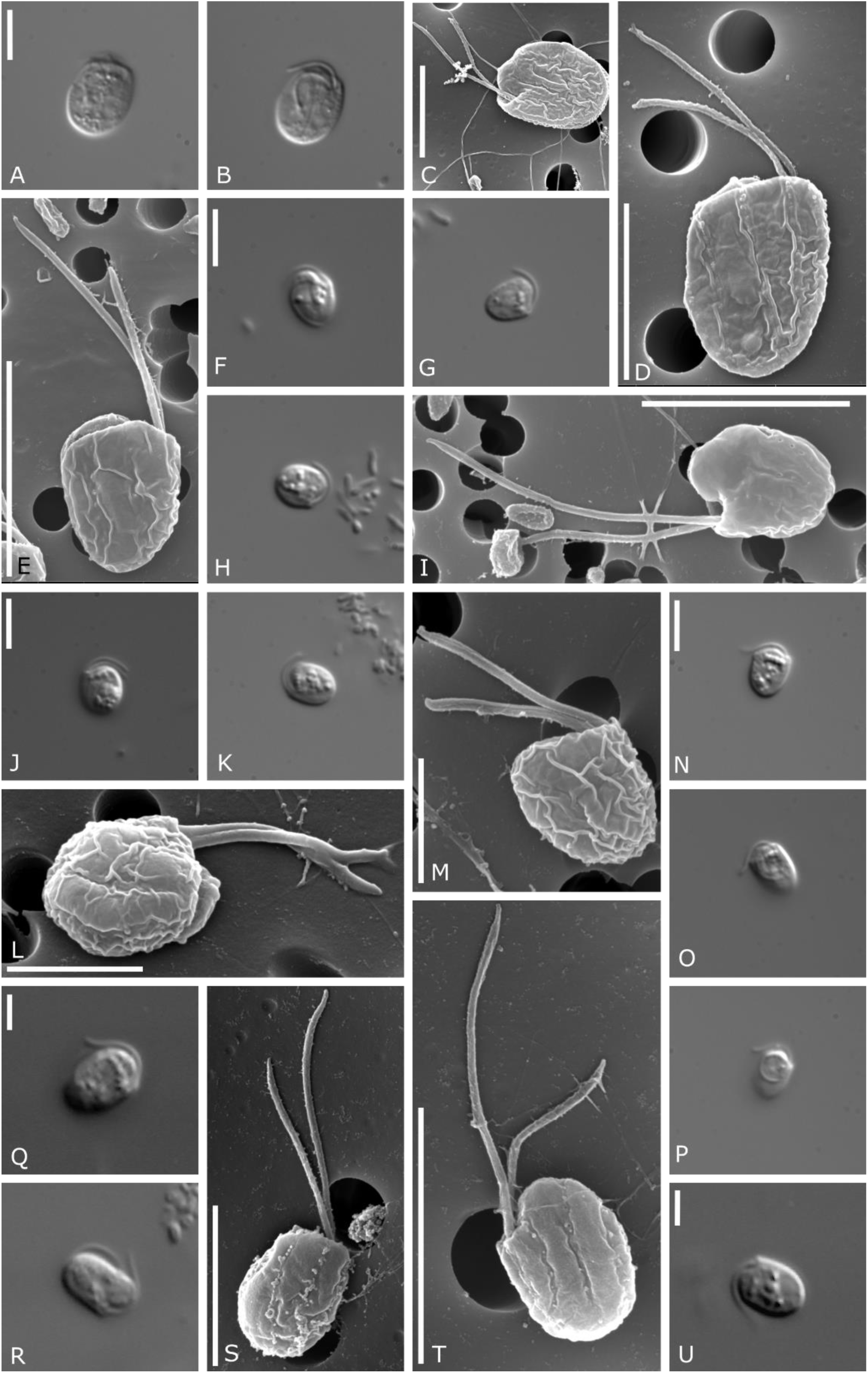
Light and electron micrographs showing: A-D *Baltigoniomonas juergensii* n. g. et n. sp. (bar indicates A, D-5 µm), E-I *Marigoniomonas* sp. – HFCC 841 (bar indicates: E, F, I-5 µm), J-M *Marigoniomonas* sp. – HFCC 272 (bar indicates: J - 5 µm, L, M-2 µm), N-P *Thalassogoniomonas amphinema* n. comb. (bar indicates: N – 5µm), Q-U *Poseidogoniomonas* sp. (HFCC 157, bar indicates: Q, U - 2µm, S - 4 µm, T - 5 µm)

Another genetically distinct strain, HFCC 1666, was placed into the newly described genus *Thalassogoniomonas* n. g. (Figure 5, N-P). Cells were obovate to cup shaped, with two unequally long flagella, the posterior carried over the cell body. The genus *Poseidogoniomonas* so far comprises two new isolates. One from our own study (HFCC157, designated *Poseidogoniomonas* sp., Figure 5, Q-U), and *P*. *duplex* n. comb. (Phanprasert et al. 2025 emend.). Both strains have cells of about 4-6 µm length and 3-4 µm width and carry two flagella, of which the shorter anterior flagellum carries fine hairs and the longer posterior flagellum is smooth. Strains differ in the number of periplast plates.

### Phylogenetic analysis

The analysis of the partial 18S rDNA data shows a clear separation between the freshwater and marine clades. The freshwater cluster is formed by the genera *Goniomonas* and the newly described genera *Limnogoniomonas* n. g. and *Aquagoniomonas* n. g. The marine cluster is formed by *Marigoniomonas* n. g., *Thalassogoniomonas* n. g. *Poseidogoniomonas* n. g. and *Cosmogoniomonas* n. g., and there is one genus, up to now only recorded from brackish water, *Baltigoniomonas* n. g. (Figure 6), closely related to *Neptunogoniomonas* n. g.

**Figure 6.**
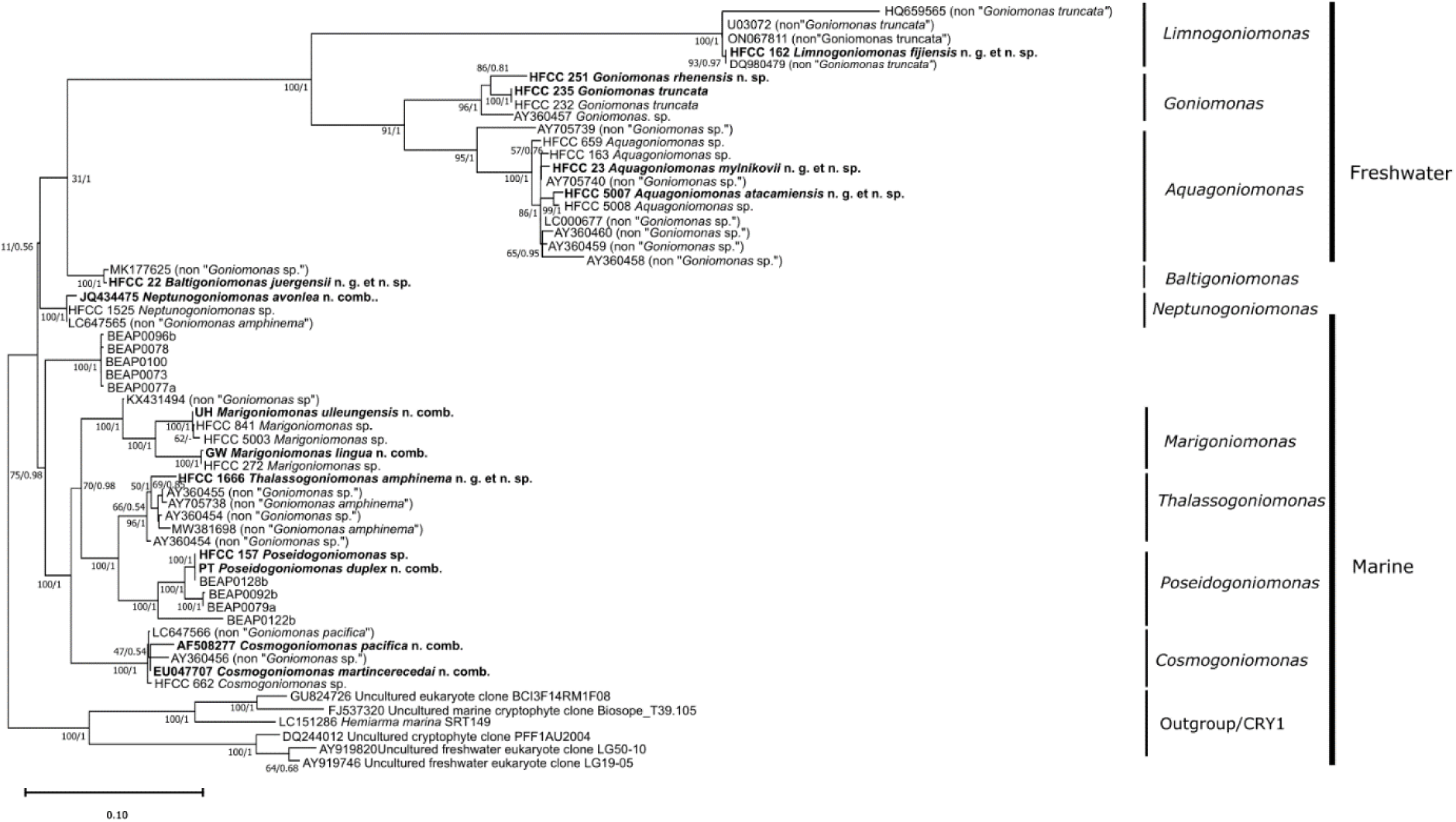
Maximum likelihood (ML) phylogenetic tree based on sequences of 18S rDNA. Investigated strains are marked in bold. Numbers indicate bootstrap values of Maximum Likelihood analysis as well as Bayesian analysis for the branches, “- “indicates a differing topology for Bayesian analysis. The unnamed sequence cluster containing sequences with the label BEAP will be discussed in further studies. Within this cluster all sequences belong to strains isolated from the Mediterranean Ocean.

Among the newly isolated and analysed strains, two cluster within the freshwater clade (HFCC 251, 235) of *Goniomonas*, one in the freshwater clade of *Limnogoniomonas* n. g. (HFCC 162), and two in the freshwater clade of *Aquagoniomonas* n. g. (HFCC 23 and 5007). Among the marine isolates one clusters in the clade *Baltigoniomonas* n. g. (HFCC 22), two in the clade *Marigoniomonas* n. g. (HFCC 841 and 272) with two from the study of Phanprasert et al. (2025), one in the clade *Thalassogoniomonas* n. g. (HFCC 1666) and one in the clade *Poseidogoniomonas* n. g. (HFCC 157). In many cases, Bayesian analysis produced stronger support of the clades than Maximum Likelihood analysis and weakly supported clades can be found both in freshwater and marine groups.

Among the freshwater clades, the neotype of *Goniomonas truncata* clusters together with two other sequences isolated from the River Rhine. A pairwise comparison revealed 0.0% difference to strain HFCC 232 and 3.8% difference to strain HFCC 251 indicating the existence of two distinct species in this cluster, strongly supported by bootstrap values (Suppl. Table 2). Sequences designated as *Goniomonas* share at least eight unique motifs within conserved regions of the 18S rDNA, with overall only two nucleotides difference in the first and fourth motif within the clade. Both differences occur in sequence AY360457. Sequence motifs in the *Goniomonas* clade share most characters with the *Aquagoniomonas* clade, followed by *Limnogoniomonas*. More dissimilarities were found in comparison to the marine and brackish water clades.

The large clade of the newly described genus *Aquagoniomonas* n. g. houses eleven sequences from strains originating from different freshwater environments. The newly described species, *Aquagoniomonas mylnikovii* n. g. et n. sp. isolated from a pond in Borok, Russia, is closely related, though not identical, to a sequence from GenBank (LC000677), and it clusters closely together with a another GenBank sequence from an UK freshwater habitat (AY705740) and a strain from our own collection (HFCC 163) isolated from the river Rhine with a relatively small genetic distance (Suppl. Table 2). The clade consists of several undescribed and undetermined strains from various geographic origins (originally assigned to *Goniomonas* despite their large genetic distances). Within this clade two strains from our own collection appear, both originating from the Rio Loa in the Atacama Desert in Chile. Strain HFCC 5007 is closely related to strain 5008. Within this clade itself, genetic differences can be high, up to 9.7 % (Suppl. Table 2). Basal branching of *Aquagoniomonas* as a sister clade to *Goniomonas* is well supported by bootstrap values (>90/0.9), while support within the large clade is partly weaker. Future studies might further resolve the phylogenetic relationship of *Aquagoniomonas* (Figure 6).

In the clade *Limnogoniomonas* n. g., strain HFCC 162, isolated from a pond on the island Fiji, clusters closely together with two sequences formerly designated as *Goniomonas truncata* (Suppl. Table 2). The genus *Limnogoniomonas* clusters as a sister clade to the other freshwater taxa is strongly supported by bootstrap values (>90/0.9) and there are no dissimilarities within the chosen motifs within this clade. While it shares some similarities with the other freshwater clades, it lacks a number of longer inserts present in the others, that leads to the individual branch. The sequence HQ659565 displays a longer branch characterized by a minor number of shorter inserts, that the other *Limnogoniomonas* n. g. do not possess (Figure 6).

The marine species clustered more separate from each other, with a shorter branch than isolates from freshwater habitats. Strain HFCC 22 from brackish environment (Baltic Sea) clusters together with a GenBank sequence (MK177625) originating from a freshwater pond on an island in Korea with only a relatively small genetic difference (Suppl. Table 4). The branch is well supported by bootstrap values and the chosen synapomorphy motifs were completely consistent between the two sequences (Figure 7). The clade *Baltigoniomonas* n. g., with *B. juergensii* n. g. et n. sp., originating from brackish waters is separated from *Neptunogoniomonas* n. g. due to several single nucleotide differences in the synapomorphy motifs (Figure 7). *Neptunogoniomonas* n. g. includes the most recently described “*G.*” *avonlea* (now *Neptunogoniomonas avonlea* n. comb.) isolated from Canadian marine waters and two other sequences, one originating from our own collection (HFCC 1525, isolated by Sabine Schiwitza from Denmark coastal waters) and one sequence that had originally been assigned to “*G. amphinema*”, LC647565. On the level of SSU rDNA, HFCC 1525 has differences to “*G*.” *avonlea* (JQ434475; Suppl. Table 5), but is identical in nuclear LSU rDNA (not included in this study). We also added several sequences (BEAP – **B**iology and **E**cology of **A**bundant **P**rotists Lab, Institute of Evolutionary Biology, CSIC-Universitat Pompeu Fabra, Barcelona) originating from organisms sampled in the Mediterranean Ocean kindly provided by Daniel Richter and Daryna Zavadska and their group to our phylogenetic analysis. Of those sequences, five formed an individual cluster supported with high bootstrap support. These potentially new species and a potentially new genus have to be described in future studies when morphological data are available.

**Figure 7.**
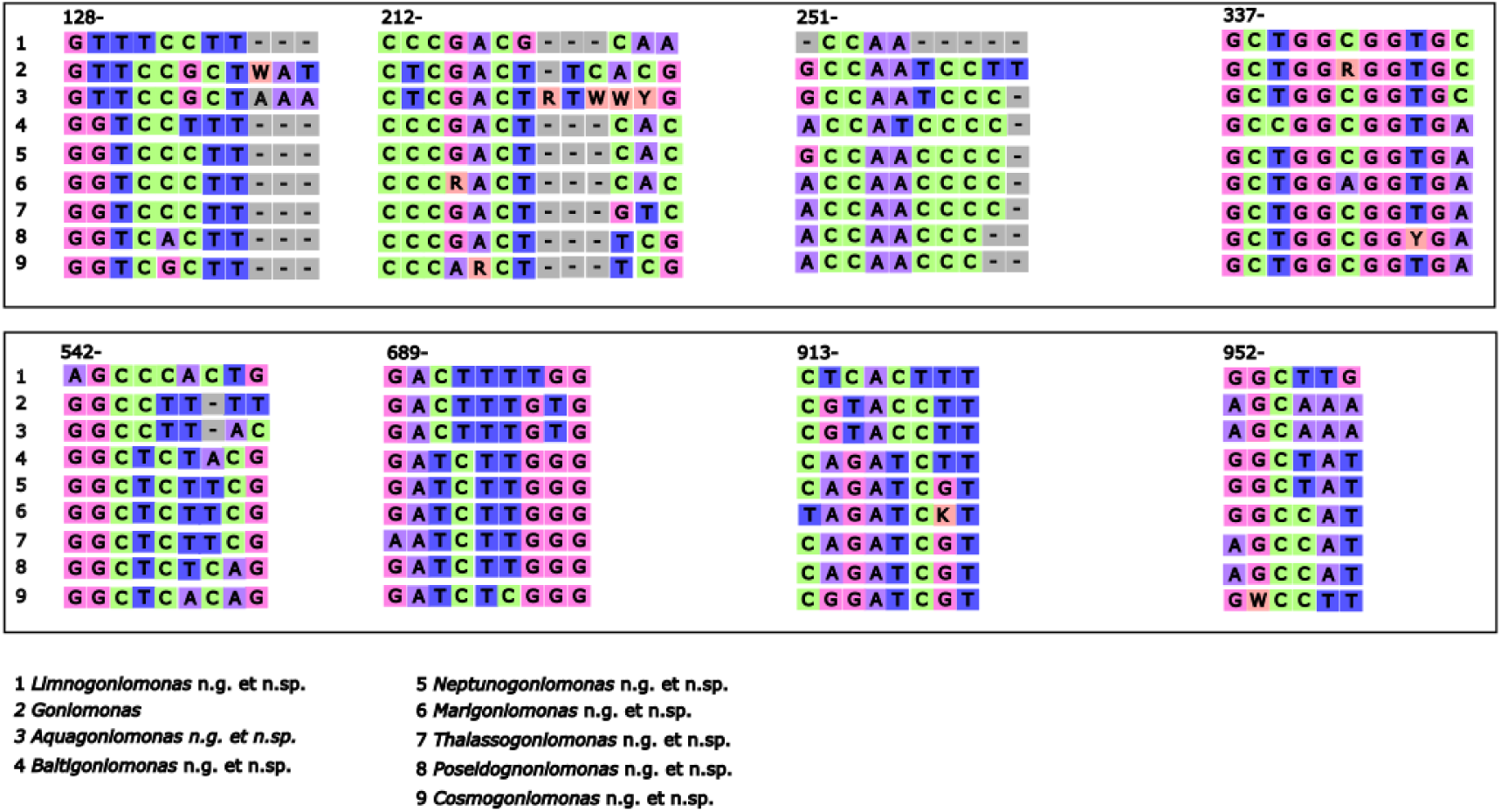
Diagram illustrating synapomorphic motifs within the 18S rDNA across nine clades, highlighting dissimilarities between the sequence clusters. The numbers above each motif represent the start position of the motif within the original alignment of this study (supplementary material). Variations in the motifs underscore evolutionary relationships and distinctions.

The remaining marine clades overlap regarding their synapomorphic motifs, though each clade has its separate identity (Figure 7). The large morphological and genetic variety of the additional marine species is structured into four clades.

Another clade with strong bootstrap support belongs to the newly created genus *Marigoniomonas* n. g. It comprises the closely related strains HFCC 841 and 5003 (Suppl. Table 7). Both originated from the Pacific at the coast of the Atacama Desert in Chile and are highly similar on the basis of the SSU rDNA. The strain HFCC 272 from the North Atlantic clusters besides the two sequences from the Pacific with a distance to strain HFCC 841. The first motif must be excluded from analysis, as it is only present in HFCC841, due to sequence length of the two other (Figure 7, in brackets). In two other motifs (two and seven), it is HFCC272 that shows a total of two single nucleotides difference underlining its separate position in the cluster. The two species *Marigoniomonas ulleungensis* n. comb. and *M. lingua* n. comb recently described by Phanprasert et al. (2025) fall into this cluster and were transferred to this new genus.

Six other sequences from marine sites form the clade designated *Thalassogoniomonas* n. g. (Suppl. Table 7). *Poseidogoniomonas* n. g. is a marine clade comprising six sequences of which one belongs to our own collection, one to the recent study of Phanprasert et al. (2025) and the other four originate from the Mediterranean Ocean (Richter and Zavadska, pers. communic., see above). The newly emended strain *P. duplex* n. comb. (former *Goniomonas duplex*, Phanprasert et al. 2025) serves as a type species for this clade. The genetic diversity in the clade is relatively large (Suppl. Table 7), still, this part of the tree is robust based on bootstrap values and we could only detect one nucleotide difference between the sequences within the chosen motifs. In motif four, strains HFCC157, *P. duplex* n. comb. and BEAP128b both display a thymine (T) at position 345 where the other two have a cytosine (C). The strain HFCC 1666 was designated as the type species of this clade and will serve as a neotype for “*Goniomonas” amphinema*.

A potentially basal genus is a clade consisting of sequences originally either named “*Goniomonas* (aff.) *amphinema*” or “*Goniomonas*” *pacifica* (*Cosmogoniomonas martincerecedai* n. comb., resp. *C. pacifica*), yet none of the sequences could serve as a type for those species. The clade is enlarged by a strain originating from our own collection, HFCC 662, isolated from the Maldives, with only minor genetic distances on the SSU rDNA level to strain LC674566 that was originally assigned to “*G*.” *pacifica*. This sequence is also very close to the sequence defined as “*G*. aff. *amphinema*” (EU047707) with only two nucleotides being different (Suppl. Table 6).

The sister group relationship of *Thalassogoniomonas, Poseidogoniomonas, Cosmogoniomonas* and *Marigoniomonas* is robust and supported by the chosen synapomorphy motifs (Figure 7).

## DISCUSSION

### Divergence of freshwater and marine strains and new genera

During this study we investigated ten different cryptophyte strains with a *Goniomonas*-like morphology. Apart from designating a valid neotype for *G. truncata*, we split the taxonomically not well resolved genus into three freshwater and six marine/brackish water genera and described six new species based on morphological and/or phylogenetic findings.

A possible splitting of the genus *Goniomonas* into further clades has already been proposed in earlier studies (Kim and Archibald, 2013; Von der Heyden et al., 2004). Von der Heyden et al. (2004) compared the branching depth between freshwater and marine species to be similar to what separates most of the prymnesiophyte genera. The phylogenetic results, and the robust values at the basal split between those two groups support the demand to especially separate those two groups. We found the largest genetic difference (38% on basis of the 18S rDNA) between the newly defined neotype of *Goniomonas truncata* (strain HFCC 235) and *Poseidogoniomonas* sp. (strain HFCC 157), a strain from one newly described marine genus.

But even between an established species like the formerly known “*G*. (aff.) *amphinema”* (EU047707, now *Cosmogoniomonas amphinema* n. comb.) and *G. truncata* we found a genetic difference of 21%. The formerly known “*G. amphinema”*, “*G. pacifica”* and “*G. avonlea”* were therefore sorted to different genera, *Thalassogoniomonas* n. g., *Cosmogoniomonas* n. g. resp. *Neptunogoniomonas* n. g. Earlier studies focussing on the 18S rDNA phylogeny of goniomonads (Deane et al., 2002; Von der Heyden et al., 2004) showed that the major difference between the freshwater and marine clades is based on several inserts which some freshwater species possess and which are absent in marine goniomonads. This pattern is confirmed by our newly described species. Deane et al. (2002) and Von der Heyden et al. (2004) found also that one sequence (UO3072) from freshwater is lacking the insert typical for all other freshwater goniomonads. The newly described genus *Limnogoniomonas* n. g. includes this specific sequence. Additionally, all other sequences that appear in the *Limnogoniomonas*-clade are lacking this insert, resulting in an individual branch, apart from the other freshwater genera. This is also reflected by the performed synapomorphy analysis. In the first motif, only *Goniomonas* and *Aquagoniomonas* show the full motif, whereas all other clusters, including *Limnogoniomonas*, display a gap after position 135. The well supported branching (as well as the large genetic distance) of the type sequence HFCC 162 to the new neotype of *G. truncata* (HFCC 235 may validate the erection of a new genus *Limnogoniomonas* n. g. The clade does also house a sequence (ON067811) originating from a study focussing on cryptic and aplastic cryptophytes (Šimek et al., 2022). The study designates, but not scientifically describes the strain as *Pseudogoniomonas* and sorts it as a sister clade to the aplastidic CRY1 lineage (Shiratori and Ishida, 2016; Shalchian-Tabrizi et al., 2008) and *Goniomonas*. However, the sequence was deposited in GenBank not under the name *Pseudogoniomonas* but *Goniomonas truncata*. Although this is initially misleading, it ultimately corresponds to our argument above, since the study separates *Pseudogoniomonas* from *Goniomonas* as we do here for *Limnogoniomonas* n. g.

Apart from the divergence between freshwater and marine/brackish strains, there was also a remarkable genetic diversity even within the different genera. In our study, the *Goniomonas* freshwater clade houses four sequences, including one of a newly described species. Between the neotype and *Goniomonas rhenensis* n. sp. we calculated a pairwise distance on basis of the 18S rDNA of approx. 3.8%. For species in the genus *Aquagoniomonas* n. g. a pairwise distance to 6% (between sequences of HFCC659 and AY360458) and up to 9.67 % (between sequences AY705739 and AY360458) were recorded. Within the marine genera, we found the following maximum genetic differences with regard to the 18S rDNA gene: *Baltigoniomonas* n. g. 0.31 %, *Neptunogoniomonas* n. g. 0.41 %, *Marigoniomonas* n. g. 3.4 %, *Thalassogoniomonas* n. g. 1.76 % and for *Poseidogoniomonas* n. g. 3.4 % (Supplementary Tables 4-7).

Some flagellate isolates originating from rather remote regions (e.g., *Thalassogoniomonas amphinema* n. comb. from Helgoland) form an individual branch. This may be signs for an allopatric speciation in remote areas/islands (Hernandez-Hernandez et al., 2021) as it was postulated for percolomonads and other protist species (Arndt et al, 2020; Hohlfeld et al., 2022). Yet, genetic differences between different protist genera can widely vary depending on the genus and family. It is therefore difficult to set up boundaries or thresholds up to where a sequence does still belong to a genus or not. We consider our separation reasonable, but are aware that further studies may change the picture again. By looking at the synapomorphy analysis, there are several differences in motifs between freshwater and marine clades and also between the different marine clades. The topology of the freshwater clades is overall more robust which could stem from the larger number of marine sequences (30 vs. 20 in the freshwater clade), reducing support. The broader topology of the tree is partly well supported which suggests that the tree correctly reflects the evolutionary relationships at higher taxonomic levels and supports our decision to split the genera and to hopefully clarify the systematic position of strains and sequences in future. Nevertheless, some positions in the tree remain uncertain, as the branching of the *Baltigoniomonas* and *Neptunogoniomonas* clade to each other as well as to the freshwater and marine clusters, highlighting their position in-between the two. The lower bootstrap values within some of the clusters could be attributed to the limited resolution of the 18S marker and could be refined with further multigene resolution. Unfortunately, only very few sequences of other marker genes are available for goniomonads at the moment in public data bases that could potentially have resolved some uncertainties.

The observed deep sequence divergence stands in contrast to the rather conserved morphological features of the established genera that does hardly allow such a clear differentiation on the first glance based on their morphology. Our study shows that features like cell size and even position and length of flagella that have been used to distinguish e.g. the former species *Goniomonas pacifica* and *G. amphinema* from each other, have lost their discriminative power as these characters are also found in other, genetically distinct species. The position, direction and length of flagella as described for “*G*. aff.” *amphinema* (Martin-Cereceda et al., 2010) can also be found in species like *Baltigoniomonas juergensii* n. g. et n. sp. and *Poseidogoniomonas* sp. (strain HFCC 157), that have at least 6% pairwise distance in the 18S rDNA to the former species. Earlier studies have already noted that even without a close link to 18S rDNA data, different genera of Cryptophyta can show surprisingly similar morphologic features (Deane et al., 2002). Hence, it is necessary to lay a stronger focus on molecular data to define species in this phylogenetic group of organisms and to use morphological features as supporting information. Still, we can observe morphological similarities within the newly described genera.

### Designation of a neotype for *Goniomonas truncata*

In public databases around 25-30 sequences are available for phylogenetic analyses of the genera investigated in this study (on basis of the 18S rDNA). After our phylogenetic analyses, four of these sequences fall into the *Goniomonas* clade. Since its first description (Fresenius, 1858) the species *Goniomonas truncata* has been analysed in multiple studies (Bernard et al., 2000; Ekelund and Patterson, 1997; Hill, 1991; Kugrens and Lee, 1991; Patterson and Lee, 2000; Vørs, 1993), but every time without any genetic material to underpin its taxonomic position. Together with other studies (von der Heyden et al., 2004), our studies show how genetically diverse the genus actually is. It is therefore hard to judge, whether the publications listed above really analysed *G. truncata* itself, or lumped multiple species into one ever-increasing description. Kim and Archibald (2013) have completed a helpful table, showing how many studies actually described each species of the genus.

The designation of strain HFCC 235 as the neotype for *Goniomonas truncata* was done, as the morphology of this strain resembles the original description and the sampling origin is close to the area where the original studies have been carried out by Fresenius. Stein (1878), placed the species from *Monas truncata* into the new genus *Goniomonas truncata*. The original description by Fresenius (1858) lists the following morphological features: cells are found in freshwater, are colourless, oval to roundish and 6-10 µm in length, cells are laterally compressed. In addition, the anterior end is truncated and carries two flagella that originate on the side and have a length of about the total cell length. Close to the anterior margin lies a band of a narrow structure (today we know that these are ejectosomes) and close to this band a contractile vacuole. The swimming behaviour is described as trembling with a frequent change in direction. The strain HFCC 235 fits well to this original description. With a cell length of 5.9–9.3 µm, an ovoid cell shape and two flagella with a similar length of 4.5–6.3 µm respectively 4.0–5.6 µm. Depending on the individual cell, the characters fall within the description given by Fresenius (1858). It also has a band of ejectosomes on the anterior margin of the cell. Additionally, HFCC 235 comes from a freshwater environment and was sampled in Germany, which is assumingly the sampling origin of both Freseniuś and Steinś strains. One could argue that Fresenius gave a rather unspecific description of a goniomonad and that, apart from the length of flagella in ratio to the cell size, his descriptions are quite universal for this genus. Nonetheless, we find this strain appropriate due to the reasons mentioned above and also on the basis of the illustrations that both Fresenius (1858) and Stein (1878) provided.

### Combining phylogenetic and morphological data

With *Limnogoniomonas* n. g. we named a clade currently consisting of 4 strains with *Limnogoniomonas fijiensis* n. g. et n. sp. as the type species. *Limnogoniomonas fijiensis* n. g. et n. sp. differs from *Goniomonas truncata* by having a larger cell length of 7.6 - 11.4 µm but being similar in width, making the cells much more elongated than the ones of *G. truncata*. In addition, *Limnogoniomonas fijiensis* n. g. et n. sp. has longer flagella than all *Goniomonas* species described within this study. In the synapomorphy motifs, the clade of *Limnogoniomonas* n. g. and *Goniomonas* differ by at least one nucleotide per motif. Their relation to each other is robustly supported by the tree topology (Figures 6 and 7).

Within the freshwater clades, three isolates from the river Rhine cluster closely together (HFCC 251, 232 and 235). Another one (HFCC 163) appears quite far away and is part of the *Aquagoniomonas* n. g. clade. HFCC 235 was designated as the neotype for the *G. truncata* and HFCC 251 was described as the new species *G. rhenensis* n. sp. With a pairwise distance of 3.8 %, the type species branches closely, though poorly supported. *G. rhenensis* n. sp. displays larger (both in length and width) cells like *G. truncata*, while the anterior and posterior flagella are of similar length. *G. rhenensis* n. sp. carries six irregularly shaped periplast plates on the right side and four on the left side, while *G. truncata* carries five left and respectively six right side periplast plates. HFCC 232 has an identical sequence to HFCC 235 and can be considered as a re-isolation of *G. truncata* from the neotype locality. The chosen motifs on basis of 18S rDNA revealed robust relationship between the *Goniomonas* species, additionally supported with the occurrence of the above-mentioned inserts, that they share only with strains of *Aquagoniomonas* n. g.

The strain HFCC 23, isolated near Borok, Russia, was described as the new species *Aquagoniomonas mylnikovii* n. g. et n. sp. It serves as the type species for this new genus. On basis of the 18S rDNA it shows a pairwise distance of 0.47 % to its closest relatives, the two undescribed sequences LC000677 and AY705460 isolated in Japan and UK, respectively. In distinction to *Goniomonas*, *Aquagoniomonas mylnikovii* n. g. et n. sp. has a similar cell length but is wider and has a longer posterior flagellum. The cell size is smaller than of *Goniomonas rhenensis* n. sp. and it has a shorter anterior and posterior flagellum than the latter. It carries four periplast plates on the left and five on the right side. HFCC 5007 was designated as *Aquagoniomonas atacamiensis* n. g. et n. sp. and has its type locality in the Rio Loa in Chile. This strain has a pairwise distance of 10.4 % to *G*. *truncata* and 0.61 % to its closest relative, a strain also originating from the Rio Loa, but was found in the mouth area of the river. *Aquagoniomonas atacamiensis* n. g. et n. sp. is of similar cell size as the type species *Aquagoniomonas mylnikovii* n. g. et n. sp. but carries longer flagella. Both species have four periplast plates on the left and five on the right side. The inner topology of the closer related sequences is mostly well supported, and except for motif two, where there is a higher level auf dissimilarities between the sequences/strains, the other motifs are completely identical within the *Aquagoniomonas* clade.

The genera *Neptunogoniomonas* n. g. and *Baltigoniomonas* n. g. comprise three sequences from public databases and two sequences from our own collection. Phylogenetically, the genus is subdivided into two branches, that we consider as different genera, as their relationship is only poorly supported in the tree. One branch contains two species from GenBank, including the species formerly known as *Goniomonas avonlea*, now *Neptunogoniomonas avonlea* n. comb. One sequence originating from Denmark just at the transition zone between North Sea and Baltic (HFCC 1525) and one sequence with marine origin (LC647565). The other branch contains strain HFCC 22, originating from the Baltic Sea, described here as *Baltigoniomonas juergensii* n. g. et n. sp. It branches together with a sequence from a freshwater pond in Korea (MK177625). Thus, the broader branch consists of strains from brackish, marine and freshwater environment. This is quite unusual as all other sequences in the lower half of the phylogenetic tree (Figure 6) have a strictly marine background. Aside from this fact, the morphology of *Baltigoniomonas juergensii* n. g. et n. sp. slightly differs from the one of *N*. a*vonlea* n. comb. With a cell length of 5.9-9.3 µm and cell width of 4.8-7.1 µm it is smaller than the latter, but the ratio of flagella length compared to the cell length seems to be similar. Both species carry mastigonemes on one flagellum whereas the other one appears to be smooth. Both species carry four rectangular periplast plates on the left side and three on the right side. The sequences show a pairwise distance of 3.4 % to each other (on basis of 18S rDNA). The synapomorphy analysis revealed several similarities between the two genera but also enough differences to justify the split. The motifs for the two clusters are completely identical within the clades.

The strains HFCC 841 and HFCC 5003 form an individual branch together with another sequence from our own collection, HFCC 272, two recently described species from South Korean sandy beaches (Phanprasert et al., 2025) and a more distant sequence from a public database (KX431494). Together they form the new genus *Marigoniomonas* n. g., with *Marigoniomonas ulleungensis* n. comb. being designated as the type species. Interestingly, *M*. *ulleungensis* n. comb. branches closely together with our strains HFCC 841 and HFCC 5003, all strains originated from coastal waters of the Pacific Ocean. On basis of the compared sequences, our isolate HFCC 5003 had only two nucleotide differences, however, at our present stage of knowledge, we cannot decide whether this is a character discriminating different species. To the next sequence in the branch, HFCC 272 and *M. lingua* n. comb., have a difference of about 3.5 %. Our strain HFCC 272 is morphologically similar to *M. lingua* n. comb., though not identical regarding the 18S rDNA (≤7 nucleotide differences). Due to methodological problems, we were not able to carry out SEM studies of strain HFCC 272. At least, the two strains are of similar size. Whether or not these isolates belong to the same species has to be investigated in future studies.

The sister clade to *Marigoniomonas* n. g. houses a variety of sequences from all over the world. Some were designated as “*G. amphinema”*, assumingly on basis of morphological observations, even though they appear genetically quite distant from the clade where “*G*. aff. *amphinema”* (EU047707) appears. Outside of this clade branches the newly designated HFCC 1666, *Thalassogoniomonas amphinema* n. comb. It is closest related to the sequence AY360454. There are no other described species which could be analysed for comparison within this clade. As the strain seems morphologically quite similar to the original description of “*G.” amphinema* (Larsen and Patterson, 1990), it was decided to use it <u>as</u> a basionym for *T. amphinema* n. comb.

The strain HFCC 157 *Poseidogoniomonas* sp. originated from the Azores islands. It clusters together with four sequences from the Mediterranean Sea (that were added for higher resolution) and one from the study of Phanprasert et al. (2025), *P. duplex* n. comb., that we designated as type for this genus. The sequences differ in 7 nucleotides, mainly coming from a small insert, that only HFCC 157 possesses (total length of alignment – 2188 nucleotides). The two new strains share several morphological characteristics, as cell size and flagella appendages, but seem to differ in the number of periplast plates.

The sixth marine clade consists of a mixture of sequences either traditionally assigned as “*G*.” (aff.) *amphinema* or “*G*. *pacifica”*. Both species had been morphologically well described in several publications (Al-Qassab et al., 2002; Ekelund and Patterson, 1997; Larsen and Patterson, 1990; Martin-Cereceda et al., 2010; Patterson and Lee, 2000; Vørs, 1993). Although our genetic analyses revealed that this clade exhibits a large genetic diversity, its members shared a high degree of morphological similarity as well as a high degree of identity in the synapomorphy motifs, where only one nucleotide difference could be detected. By defining “*G*. aff. *amphinema*” and the associated sequence EU047707 as *Cosmogoniomonas martincerecedai* n. comb. as the type species, we tried to establish a solid foundation for the taxonomy. The comprehensive species description by Martin-Cereceda et al. (2010) serves as an ideal reference for comparing this species with others. “*G*. *pacifica”*, unfortunately, lacks ultrastructural descriptions. Within our phylogenetic analysis, two sequences were assigned to “*G*.” *pacifica.* With a pairwise distance of 0.12%, one of these sequences (LC647566) shows only minor pairwise differences to *Cosmogoniomonas martincerecedai* n. comb. (EU047707) and it is likely that it has been falsely assigned. Consequently, we chose to redefine the remaining sequence (AF508277) as a reference for *Cosmogoniomonas pacifica* n. comb. as it originates from Australia, and was sampled from the Pacific Ocean.

Our study, as other studies before, demonstrates the large diversity within the genus *Goniomonas* and offers a solution by splitting the genus into several individual genera. Still, we are aware that this might not be the last taxonomic revision that will be made for goniomonads. The large intraspecific differences hint at a much larger diversity and, by adding more sequences, may allow an even finer resolution (and splitting) either on the genus or species level. As already mentioned, it is hard to draw a line on basis of genetic differences, especially when the morphological bauplan is similar. Whereas differences within the marine genera ranged around 4 %, they were even higher within freshwater genera such as *Aquagoniomonas* n. g.

As with many protist groups, the diversity of goniomonads is still largely unknown, but, particularly with the help of modern sequencing techniques, we are slowly taking steps to get an idea about their diversity. Nevertheless, modern taxonomy, which complements molecular databases, is the backbone for all metabarcoding studies. It is crucial to keep the taxonomy up to date, otherwise much of the diversity will be lost or remain cryptic. As more and more sequences are generated, it is important to critically examine previous taxonomic decisions and revise them if necessary.

## TAXONOMIC SUMMARY

Assignment: Eukaryota, Cryptista, Cryptophyta, Goniomonadea, Goniomonadida

## TAXONOMY OF NOVEL GENERA AND SPECIES

### *Goniomonas* Stein 1878, n. emend. Sachs et Arndt

#### Diagnosis

A free living, colourless, bi-flagellated heterotrophic cryptophyte only found in freshwater habitats, bacterivorous. Cells are laterally compressed, slightly longer than wide, ovoid to pear shaped. Anterior end slightly angled, posterior end rounded. Cells carry two flagella of subequal length that can carry fine hairs or appendages. The cell body is covered by irregularly shaped, elongated periplast plates.

Type species. *G. truncata*

### *Goniomonas truncata* Stein 1878, emend. Sachs et Arndt

#### Diagnosis

Free living, colourless, bi-flagellated cryptophyte from freshwater environment. Laterally compressed cells, slightly elongated, ovoid to pear-shaped cell body, rounded towards posterior end, tapered, angularly truncated anterior end, with 5.9 – 9.3 µm length (average 7.3 µm, SD=1.0, n=22) and 4.4–7.1 µm width (average 5.5 µm, SD = 0.6, n = 22) of cell body. Two flagella of subequal length arise from an anterior groove and are both directed anterior-laterally with the anterior flagellum of 4.5–6.3 µm (average 5.4, SD=0.6, n=17) length and the posterior flagellum being 4.0–5.6 µm (average 4.9, SD=0.5, n=16) long. Both flagella are equipped with fine hairs, the shorter flagellum carries additional spikey appendages. Cells carry five irregularly shaped periplast plates on the left side and six on the right side. A band of ejectosomes is visible on the anterior margin of the cell with 1.7–2.8 µm (average 2.3, SD=0.4, n=7) in length. Between the junctions of the periplast plates, small ejectosomes arise as globule like structures. Additionally, species of the genus *Goniomonas* display several unique motifs within sequences of the 18S rDNA (Figure 7) that are distinct from other genera.

#### Neotype location

Strain HFCC 235 was obtained from the River Rhine, Cologne, 50°49’34"N 6°35’34"E

#### Neotype material

The type culture is currently deposited in the Heterotrophic Flagellate Collection Cologne, as strain HFCC 235, specimen shown in Figure 1, A. SEM filters are deposited at the Biology Centre of the Museum of Natural History in Upper Austria, Linz, at number XXX. Deep-frozen material is deposited at the Heterotrophic Flagellate Culture Collection Cologne.

#### Gene sequence data

The 18S rDNA sequence has been deposited at GenBank with the Accession Number xxx

#### ZooBank ID

XXX (will be provided prior to publication)

### *Goniomonas rhenensis* n. sp. Sachs, Nitsche et Arndt

#### Diagnosis

Free living, colourless, bi-flagellated cryptophyte from freshwater environment. Laterally compressed cells, slightly elongated, ovoid to pear-shaped cell body, rounded towards posterior end, tapered, angularly truncated anterior end, with 7.8–10.8 µm (average 9.3 µm, SD = 0.4, n =20) length and 5.3–7.9 µm (average 6.4 µm, SD = 0.8, n = 20) width of cell body. Two flagella of subequal length arise from an anterior groove and are directed anterior-laterally with the anterior flagellum of 3.9–6.7 µm (average 5.8 µm, SD=0.7, n = 20) length and the posterior flagellum 3.2–5.5 µm (average 4.6 µm, SD=0.5, n =20) long. Cells carry six irregularly shaped periplast plates on the right side and four on the left side. A slim band of ejectosomes is visible on the anterior margin of the cell with 1.6–1.9 µm (average 1.7 µm, SD = 0.1, n = 6) in length. Small ejectosomes can be seen between the junctions of the periplast plates.

#### Type location

Strain HFCC 251 was obtained from the River Rhine, Cologne, 50°49’34"N 6°35’34"E

#### Type material

The type culture is currently deposited in the Heterotrophic Flagellate Collection Cologne, as strain HFCC 251, specimen shown in Figure 1, K. SEM filters are deposited at the Biology Centre of the Museum of Natural History in Upper Austria, Linz, at number XXX. Deep-frozen material is deposited at the Heterotrophic Flagellate Culture Collection Cologne.

#### Gene sequence data

The 18S rDNA sequence has been deposited at GenBank with the Accession Number xxx

#### ZooBank ID

XXX (will be provided prior to publication)

#### Etymology

The species name (lat. “from the Rhine”) is based on the type locality of the strain, the River Rhine

### *Aquagoniomonas* n. g. Sachs et Arndt

#### Diagnosis

A free living, colourless, bi-flagellated heterotrophic cryptophyte from freshwater habitats, bacterivorous. Cell body laterally compressed, elongated, lean to slightly ovoid shape. Posterior end slightly tapered and rounded, proximal end truncated and angled. Cells carry two smooth flagella of subequal length, anterior flagellum slightly longer than posterior, both beeing directed anterior laterally. The cell body is covered with irregularly shaped, elongated periplast plates, four on the left side and five on the right side, with small ejectosomes arising from in between the junctions. A band of ejectosomes can be seen transversally at the anterior margin. Additionally, species of the genus *Aquagoniomonas* display several unique motifs within sequences of the 18S rDNA (Figure 7) that are distinct from other genera.

#### Type species

*Aquagoniomonas mylnikovii* n. g. et n. sp.

#### Etymology

The genus name refers to the origin of the strains in this genus, the freshwater environment (lat. "aqua” - water)

### Aquagoniomonas mylnikovii n. sp. Sachs et Arndt

#### Diagnosis

Free living, colourless, bi-flagellated cryptophyte from freshwater environment. Laterally compressed cells, elongated, lean cell body, rounded, slightly tapered towards posterior end, angularly truncated anterior end, with 6.1–9.7 µm (average 7.7 µm, SD = 1.1, n = 19) length and 4.0–5.6 µm (average 4.7, SD = 0.5, n = 20) width of cell body. Two flagella of subequal length arise from an anterior groove and are directed anterior-laterally with the anterior flagellum being slightly longer with 3.8–6.7 µm (average 5.5 µm, SD = 0.8, n = 8) length and the posterior flagellum 2.0–5.5 µm (average 4.0 µm, SD=1.2, n =8) long. Cells carry four irregularly shaped periplast plates on the left side and five on the right side. A prominent band of ejectosomes is visible on the anterior margin of the cell with 1.8–2.5 µm (average 2.0 µm, SD = 0.2, n = 8) in length, as well as small ejectosomes that arise from in between the junctions of the periplast plates.

#### Type location

Pond, Borok near Jaroslawl, Russia, 58°03’27"N 38°15’37"E

#### Type material

The type culture is currently deposited in the Heterotrophic Flagellate Collection Cologne, as strain HFCC 23, specimen shown in Figure 1, N-O. SEM filters are deposited at the Biology Centre of the Museum of Natural History in Upper Austria, Linz, at number XXX. Deep-frozen material is deposited at the Heterotrophic Flagellate Culture Collection Cologne.

#### Gene sequence data

The 18S rDNA sequence has been deposited at GenBank with the Accession Number xxx

#### ZooBank ID

XXX (will be provided prior to publication)

#### Etymology

This species is dedicated to and in remembrance of Alexander P. Mylnikov for his unparalleled contributions to protozoological research. He isolated this strain 30 years ago.

### *Aquagoniomonas atacamiensis* n. sp. Sachs, Nitsche et Arndt

#### Diagnosis

Free living, colourless, bi-flagellated cryptophyte from freshwater environment. Laterally compressed cells, slightly elongated ovoid cell body, rounded towards posterior end, angularly truncated anterior end, with 6.0–9.0 µm (average 7.7 ± 0.8 µm; n = 15) length and 3.9–5.9 µm (average 5.0 ± 0.6 µm, n = 15) width of cell body. Two flagella of subequal length arise from an anterior groove and are directed anterior-laterally with the anterior flagellum being slightly longer with 7.5–7.6 µm length (average 7.6 ± 0.1, n = 3) than the posterior flagellum being 5.6 µm (average 5.6 ± 0.0 µm, n = 3) long. Cells carry four irregularly shaped periplast plates on the left side and five on the right side. A band of ejectosomes following the angle of the anterior margin is visible and 1.3–2.4 µm (average 2.0 ± 0.5 µm, n = 4) in length as well as small ejectosomes that arise between the borders of the periplast plates.

#### Type location

Atacama, Rio Loa mouth, Chile, 21°25’41’’S, 70°03’15’’W

#### Type material

The type culture is currently deposited in the Heterotrophic Flagellate Collection Cologne, as strain HFCC 5007, specimen shown in Figure 1, T. SEM filters are deposited at the Biology Centre of the Museum of Natural History in Upper Austria, Linz, at number XXX. Deep-frozen specimens are deposited at the Heterotrophic Flagellate Culture Collection Cologne.

#### Gene sequence data

The 18S rDNA sequence has been deposited at GenBank with the Accession Number xxx

#### ZooBank ID

XXX (will be provided prior to publication)

#### Etymology

The species name refers to the geographic origin of the strain. The Rio Loa runs over 400 km through the Atacama Desert until it disembogues into the Pacific Ocean and is dedicated to the friendly people of the Atacama region.

### *Limnogoniomonas* n. g. Sachs et Arndt

#### Diagnosis

Non-photosynthetic, colourless, single celled, bacterivorous cryptophycean bi-flagellate from freshwater habitats. Cell body laterally compressed, clearly elongated, lean shape. Posterior end slightly rounded, proximal end truncated, obtuse and slightly angled. Cells carry two smooth flagella of subequal length, anterior flagellum slightly longer than posterior, both directed anterior laterally. Cell body covered with irregularly shaped, elongated periplast plates, four on the left side and five on the right side. In between those plates, small ejectosomes can be seen between the junctions of the periplast plates. At the anterior margin of the cell, a more prominent band of ejectosomes appears. Additionally, species of the genus *Limnogoniomonas* display several unique motifs within sequences of the 18S rDNA (Figure 7) that are distinct from other genera.

#### Type species

*Limnogoniomonas fijiensis* n. g. et n. sp.

#### Etymology

The genus name is derived from the fact that the clade of *Limnogoniomonas* consists of sequences only originating from freshwater samples.

#### Species included

*Limnogoniomonas fijiensis* n. g. et n. sp.

### Limnogoniomonas fijiensis n. sp. Sachs et Arndt

#### Diagnosis

Free living, colourless, bi-flagellated cryptophyte from freshwater environment. Laterally compressed cells, elongated cell body, rounded towards posterior end, angularly truncated and obtuse anterior end, with 7.6–11.5 µm with (average 9.3 ± 1.1 µm, n = 20) length and 3.9–7.3 µm (average 5.4 ± 0.9 µm, n = 20) width of cell body. Two flagella of subequal length arise from an anterior groove and are directed anterior-laterally with the anterior flagellum being slightly longer with 6.0–7.7 µm (average 6.8 ± 1.1 µm, n = 14) length than the posterior flagellum 4.5–6.6 µm (average 5.8 ± 0.5 µm, n = 13) long. Cells carry four irregularly shaped periplast plates on the left side and five on the right side. A band of ejectosomes following the angle of the anterior margin is visible with 1.7–2.7 µm (average 2.2 ± 0.4 µm, n =7) in length. Smaller, nodule like ejectosomes arise in between the periplast plates.

#### Type location

Rainforest pond, Suva, Island of Fiji, 18°08’29"S, 178°26’29"E

#### Type material

The type culture is currently deposited in the Heterotrophic Flagellate Collection Cologne, as strain HFCC 162, specimen shown in Figure 3, B. SEM filters are deposited at the Biology Centre of the Museum of Natural History in Upper Austria, Linz, at number XXX. Deep-frozen specimens are deposited at the Heterotrophic Flagellate Culture Collection Cologne.

#### Gene sequence data

The 18S rDNA sequence has been deposited at GenBank with the Accession Number xxx

#### ZooBank ID

XXX (will be provided prior to publication)

#### Etymology

The species name refers to the geographical origin of the type species, the Fiji Islands in the Pacific Ocean and is dedicated to the friendly people of the Fiji archipelago.

### *Baltigoniomonas* n. g. Sachs et Arndt

#### Diagnosis

A non-photosynthetic, colourless, single celled, bacterivorous cryptophycean bi-flagellate from marine or brackish habitats. Cell-body rounded, ovoid in shape, rounded posterior end, truncated anterior end. Cells have four periplast plates on the left side and three on the right side. Two flagella of subequal length, slightly longer than cell length and with different orientation. During movement, anterior flagellum is directed anterior-laterally, posterior flagellum is directed towards the cell. The Anterior flagellum is equipped with hairs, posterior is flagellum smooth. Ejectosomes can be seen arising between the junctions of the periplast plates. Species of the genus *Baltigoniomonas* display several unique motifs within sequences of the 18S rDNA (Figure 7) that are distinct from other genera.

#### Type species

*Baltigoniomonas juergensii* n. sp. Sachs et Arndt

#### Etymology

The prefix “*Balti*” refers to the Baltic Sea, where the type species of this genus was sampled.

### Baltigoniomonas juergensii n. sp. Sachs et Arndt

#### Diagnosis

Free living, colourless, bi-flagellated cryptophyte from brackish environment. Laterally compressed, ovoid cells with a truncated anterior end, 5.9–9.3 µm (average 7.7 ± 0.9 µm, n = 20) long and 4.8–7.1 µm (average 6.1 ± 0.6 µm, n = 20) wide. Flagella are of unequal length, the anterior flagellum is directed anterior-laterally and 4.9–7.6 µm (average 6.7 µm, SD=0.6, n = 20) long and is equipped with mastigonemes, posterior flagellum is directed towards the cell body and oftentimes carried laying over the cell and of 5.5–9.0 µm (average 7.2 ± 0.9 µm, n = 20) length and smooth. Cells carry four rectangular periplast plates on the left side and three on the right side. No band of ejectosomes could be observed at the anterior margin of the cell, but smaller nodule like ejectosomes between the periplast plates.

#### Type location

Island Hiddensee, Germany, Baltic Sea, 54°35’37"N 13°06’34"E

#### Type material

The type culture is currently deposited in the Heterotrophic Flagellate Collection Cologne, as strain HFCC 22, specimen shown in Figure 4, D. SEM filters are deposited at the Biology Centre of the Museum of Natural History in Upper Austria, Linz, at number XXX. Deep-frozen specimens are deposited at the Heterotrophic Flagellate Culture Collection Cologne.

#### Gene sequence data

The 18S rDNA sequence has been deposited at GenBank with the Accession Number xxx

#### ZooBank ID

XXX (will be provided prior to publication)

#### Etymology

The species is dedicated to Klaus Jürgens for his major contributions to the research on microbes of the Baltic Sea.

### *Neptunogoniomonas* n. g. Sachs et Arndt

#### Diagnosis

A colorless, bacterivorous marine flagellate with flattened cell body. Cell 8–11 µm long, 6–7 _m wide, somewhat truncated at the anterior. Two subequal flagella are slightly longer than the cell and arise from an anterior depression. One flagellum is directed forward whereas the other runs backward toward the cell posterior through a longitudinal groove. A conspicuous band of large ejectosomes runs across the anterior margin. Younger cells tend to have a more pointed posterior end compared to older cells, which have a more rounded end. When swimming, the cell rotates around its longitudinal axis. Swimming cells slide along surfaces with the posterior flagellum trailing beneath the cell.” (Kim and Archibald, 2013). Small ejectosomes arise between the junctions of the periplast plates. Additionally, species of the genus *Neptunogoniomonas* display several unique motifs within sequences of the 18S rDNA (Figure 7) that are distinct from other genera.

#### Type species

*Neptunogoniomonas avonlea* (Kim et Archibald 2013) n. comb. Sachs et Arndt

#### Etymology

The prefix Neptuno refers to the roman god of the oceans, with reference to the Atlantic Ocean where the type species of the genus has been sampled.

### *Marigoniomonas* n. g. Sachs et Arndt

#### Diagnosis

Non-photosynthetic, colourless, single celled, bacterivorous chryptophycean bi-flagellate from marine environments. Cells laterally compressed, roundish to d-shaped due to rounded posterior end and truncated, angled anterior end, angle pointing upwards from the attachment point of the two unequally long flagella, whereas the shorter flagellum is directed anterior laterally and the longer directed towards the cell body. Cells carry three to five periplast plates on the left side and three to four on the right side, with small ejectosomes arising in-between them. A band of ejectosomes can be seen at the anterior margin of the cell.

#### Type species

*Marigoniomonas lingua* (Phanprasert et al. 2025) n. comb. Sachs et Arndt

#### Etymology

The genus name refers to the marine environment as an origin for all included taxa

### *Marigoniomonas lingua* (Phanprasert et al. 2025) n. comb. Sachs et Arndt

#### Basionym

*Goniomonas lingua* Phanprasert et al. 2025

For diagnosis of the original description of *Marigoniomonas lingua* n. comb. see Phanprasert et al. (2025)

### *Marigoniomonas ulleungensis* (Phanprasert et al. 2025) n. comb. Sachs et Arndt

#### Basionym

*Goniomonas ulleungensis* Phanprasert et al. 2025

For diagnosis of the original description *Marigoniomonas ulleungensis* n. comb. see Phanprasert et al. (2025)

### *Thalassogoniomonas* n. g. Sachs et Arndt

#### Diagnosis

Non-photosynthetic, colourless, single celled, bacterivorous chryptophycean bi-flagellate from marine environments. Cells cup shaped, tapered posterior end and truncated, yet rounded anterior end. Two unequally long flagella arise from anterior margin of the cell, the anterior flagellum anterior-laterally directed, posterior flagellum carried over the cell in an almost curled manner. A prominent band of ejectosomes is visible transversally at the anterior margin of the cell. Species of the genus *Thalassogoniomonas* display several unique motifs within sequences of the 18S rDNA (Figure 7) that are distinct from other genera.

#### Type species

Thalassogoniomonas amphinema n. comb.

#### Etymology

The prefix “Thalasso“ (Greek: thalassa – ocean) refers to the ocean as the environment of origin.

*Thalassogoniomonas amphinema* (Larsen et Patterson, 1990) n. comb. Sachs et Arndt

##### Basionym

*Goniomonas amphinema* Larsen & Patterson, 1990

#### Diagnosis

Free-living, colourless, bi-flagellated cryptophyte from marine environment. Laterally compressed cells, cup shaped, slightly elongated, with tapered posterior and truncated, obtuse anterior end, cells of 4.3–5.7 µm (average 5.2 ± 0.5 µm, n = 19) length and 3.1–4.6 µm (average 3.8 ± 0.5 µm, n = 19) width of cell body. Two flagella of unequal length, anterior flagellum with 3.3-6.3 µm (average 5.1 ± 0.7 µm, n =16) and directed anterior-laterally, posterior flagellum generally directed towards the cell body, 5.2–8.6 µm (average 7.4 ± 0.8 µm, n = 18) long. A prominent band of ejectosomes is visible on the anterior margin of the cell with 1.7–2.3 µm (average 1.9 ± 0.2 µm, n = 12) length.

#### Neotype location

North Sea, shore of the Island Helgoland (Germany), 54°11‘01"N 7°53‘27"W from a depth of 950 m.

#### Neotype material

The neotype culture is currently deposited in the Heterotrophic Flagellate Collection Cologne, as strain HFCC 1666, specimen shown in Figure 4, N. SEM filters are deposited at the Biology Centre of the Museum of Natural History in Upper Austria, Linz, at number XXX. Deep-frozen specimens are deposited at the Heterotrophic Flagellate Culture Collection Cologne.

#### Gene sequence data

The 18S rDNA sequence has been deposited at GenBank with the Accession Number xxx

#### ZooBank ID

XXX (will be provided prior to publication)

#### Remarks

Since the light microscopic images of *G. amphinema* (Larsen and Patterson, 1990), cell size and width, as well as the length to width ratio appears to be very similar to HFCC1666, it is reasonable to designate *Thalassogoniomonas amphinema* as neotype for *Goniomonas amphinema*.

### *Poseidogoniomonas* n. g. Sachs et Arndt

#### Diagnosis

Non-photosynthetic, colourless, single celled, bacterivorous cryptophycean bi-flagellate from marine environments. Cells laterally compressed, roundish, slightly elongated cell body, sides almost parallel to each other. Cells carry three to five periplast plates on the right side and four to five on the left side. Two unequal flagella, with different directions, smooth shorter anterior flagellum directed anterior laterally and longer posterior flagellum directed towards or curved over the cell equipped with fine hairs. Ejectosomes appear as a transversal band as well as small globules in between the periplast plates. Species of the genus *Poseidogoniomonas* display several unique motifs within sequences of the 18S rDNA (Figure 7) that are distinct from other genera.

#### Type species

Poseidogoniomonas duplex n. comb.

#### Etymology

The addition “*Poseido*” refers to the Greek word for god of the ocean, Poseidon, and points to the marine origin of the type strain.

### *Poseidogoniomonas duplex* (Phanprasert et al., 2025) n. comb. Sachs et Arndt

#### Basionym

*Goniomonas duplex* Phanprasert et al., 2025

For diagnosis of the original description of *Poseidogoniomonas duplex* n. comb. see Phanprasert et al. (2025).

### *Cosmogoniomonas* n. g. Sachs et Arndt

#### Diagnosis

Non-photosynthetic, colourless, single celled, bacterivorous cryptophycean bi-flagellate from marine environments. Cells ovate with a truncated anterior and rounded posterior end, dorso-ventrally flattened. Two flagella of similar length, one pointing forward (can be equipped with appendages), one bent ventrally, trailing. Cells are covered with periplast plates (exact number not stated, see remarks). Between the plates, lines of ejectosomes can be found in the ridges. An additional transversal band of ejectosomes can be seen at the anterior end of the cell. Species of the genus *Cosmogoniomonas* display several unique motifs within sequences of the 18S rDNA (Figure 7) that are distinct from other genera.

#### Type species

*Cosmogoniomonas martincerecedai* (Martin-Cereceda et al., 2010) n. comb. Sachs et Arndt

#### Etymology

“*Cosmo*” refers to the supposedly cosmopolitan distribution of this genus.

#### Remarks

Species within the genus *Cosmogoniomonas* n. g. are morphologically relatively similar, but genetically relatively diverse within a significantly separated clade (Figure 5). Since morphological features are well described, we here define the 18S rDNA genotype EU047707 (Martin-Cereceda et al., 2010) as the neotype sequence for *Cosmogoniomonas martincerecedai* n. comb. The detailed description by Martin-Cereceda et al. (2010) gives the best reference for this species. For “*G*. *pacifica”* light microscopic studies exist (Larsen and Patterson, 1990). In GenBank, there are only two sequences at the moment that were assigned to “*G*. *pacifica”*. One sequence (LC647566) only has a minor (0.12 %) pairwise difference to *Cosmogoniomonas martincerecedai* n. comb. (EU047707). Hence, we have decided to define the other existing sequence (AF508277, Deane et al., 2002) as the neotype sequence for *Cosmogoniomonas pacifica* n. comb. as it includes the description of Larsen and Patterson (1990) in the description for this species. Our choice is underpinned by the fact that AF208277 originates from the Australian Pacific.

### *Cosmogoniomonas martincerecedai* (Martin-Cereceda et al., 2010) n. comb. Sachs et Arndt

#### Diagnosis adapted from Martin-Cereceda et al. (2010)

Free-living, colourless, bi-flagellated cryptophyte from marine environment. Cells have an anterior truncate oval shape and are 5.6 ± 0.51 µm long and 3.5 ± 0.57 µm wide (n = 90). One straight shorter flagellum beating jerkily in all directions (5.18 ± 0.44 µm long), more rapidly than the trailing, longer one (6.32 ± 0.45 µm long, n = 45). The short flagellum carries flagellar appendages. The flagellar vestibulum is about one fourth of the cell length. A line of ejectosomes appearing transversally in the anterior end of the cell. A cytopharynx structure is often visible along the longitudinal axis of the cell. Between the periplast plates, rows of emerging ejectosomes are visible. The central part of the anterior part of the cell appears raised. On the ventral anterior side, a granular area is visible.

#### Type location

Menai Strait, North Wales, UK, 53.231N, 4.164 W

#### Type material

Culture collection of Algae and Protozoa, CCAP 980/2, Figure 5 in Martin-Cereceda et al. (2010).

#### Gene sequence data

The 18S rDNA sequence has been deposited at GenBank with the Accession Number EU047707 (Martin-Cereda et al., 2010).

*Cosmogoniomonas pacifica* (Larsen et Patterson, 1990) n. comb. Sachs et Arndt

##### Basionym

Basionym: *Goniomonas pacifica* Larsen and Patterson (1990)

#### Diagnosis of the original description by Larsen and Patterson (1990)

**“**Cell outline ovate, truncate anteriorly, rounded posteriorly, 8-10 µm long, 6-8 m wide, laterally compressed. With a transverse band of ejectosomes near the anterior end. The left-hand side of the cell with 2-3 longitudinal very delicate ridges, the right-hand side with 3 ridges which are usually more distinct. Nucleus situated in the middle of the cell near the dorsal side. Two, equal or slightly unequal (dorsal longer) flagella emerge from a small depression anteriorly near the dorsal surface, about 3/4 the cell length. At their insertion the bases lie parallel to the axis of the cell. When swimming the flagella diverge; the one situated closest to the dorsal surface points forwards and bends only slightly in dorsal direction, while the other flagellum usually bends towards the ventral side. The cell skids parallel to the substratum when moving.”

#### Type location

Port Phillip Bay, Melbourne, Victoria, Australia.

#### Type material

Figure 1g in Larson and Patterson (1990).

#### Gene sequence data

The 18S rDNA sequence has been deposited at GenBank with the Accession Number AF508277 (Deane et al., 2002).

#### Remarks

Deane et al. (2002) have identified their strain as “*Goniomonas” pacifica*. Since this strain was isolated in the region from where also the originally described “*Goniomonas” pacifica* originated from, we decided to assign the sequence provided by Deane et al. (2002) to the species description. The culture of this species is deposited at the Provasoli-Guillard Center for Culture of Marine Phytoplankton, strain CCMP1869.

### CRediT authorship contribution statement

Maria Sachs: Data curation, Formal analysis, Investigation, Visualization, Writing – original draft. Frank Nitsche: Investigation, Supervision, Writing – review & editing. Hartmut Arndt: Conceptualization, Investigation, Supervision, Funding acquisition, Project administration, Writing – review & editing.

## Acknowledgements

Special thanks go to Rosita Bieg, Brigitte Gräfe and Anke Pyschny for their valuable technical support. Many thanks go also to Sabine Schiwitza, who provided strain HFCC 1525 to the study. We are also very thankful to many students helping in field work in the Atacama Desert. Especially, we would like to express our sincere thanks to Daniel Richter and Daryna Zavadska from the Institute of Evolutionary Biology (CSIC, Biology and Ecology of Abundant Protists Lab, Barcelona), for providing us with several sequences originating from organisms sampled in the Mediterranean Ocean which allowed us to stabilize the analysis of phylogenetic relationships. This research was supported by grants to H.A. from the German Federal Ministry for Education and Research (Bundesministeriumf ür Bildung und Forschung, BMBF) within its DAM pilot mission project MGF Baltic Sea (MGF-Ostsee;grant number 03F0848A), by the Deutsche Forschungsgemeinschaft (DFG, German Research Foundation) regarding the Collaborative Research Centre 1211 “Evolution at the dry limit” (project number 268236062 (SFB 1211; B03, B02) and regarding the Project AR288/23,24 and MerMet 17-11 and by a grant to F.N. (DFG, SPP 1991, project NI 1097/4).

## Data availability

Data are published or will be made available on request.

## References

Abramoff, M.D., Magalhaes, P.J. & Ram, S.J. (2004) Image processing with ImageJ. Biophotonics International, 11(7), 36–42.

Al-Qassab, S., Lee, W.J., Murray, S., Simpson, A.G.B. & Patterson, D.J. (2002) Flagellates from stromatolites and surrounding sediments in Shark Bay, Western Australia. Acta Protozoologica, 41, 91 – 144.

Archibald, J.M. (2020) Cryptomonads. Current Biology, 30(19), R1114–R1116. Available from: 10.1016/j.cub.2020.08.101

Arndt, H., Dietrich, D., Auer, B., Cleven, E., Gräfenhan, T., Weitere, M. & Mylnikov, A. (2000) Functional Diversity of Heterotrophic Flagellates in Aquatic Ecosystems. In: Leadbeater, B.S.C. & Green, J.C. (Eds.) The Flagellates: Unity, Diversity and Evolution. Taylor & Francis, CRC Press, London, pp. 240–268. Available from: 10.1201/9781482268225

Arndt, H., Ritter, B., Rybarski, A., Schiwitza, S., Dunai, T. & Nitsche, F. (2020) Mirroring the effect of geological evolution: Protist divergence in the Atacama Desert. Global and Planetary Change, 190, 103193. Available from: 10.1016/j.gloplacha.2020.103193

Bernard, C., Simpson, A.G.B. & Patterson, D.J. (2000) Some free-living flagellates (Protista) from anoxic habitats. Ophelia, 52(2), 113–142. Available from: 10.1080/00785236.1999.10409422

Carr, M., Richter, D.J., Fozouni, P., Smith, T.J., Jeuck, A., Leadbeater, B.S.C. & Nitsche, F. (2017) A six-gene phylogeny provides new insights into choanoflagellate evolution. Molecular Phylogenetics and Evolution, 107, 166–178. Available from: 10.1016/j.ympev.2016.10.011

Cenci, U., Sibbald, S.J., Curtis, B.A., Kamikawa, R., Eme, L., Moog, D., Henrissat, B., Maréchal, E., Chabi, M., Djemiel, C., Roger, A.J., Kim, E. & Archibald, J.M. (2018) Nuclear genome sequence of the plastid-lacking cryptomonad *Goniomonas avonlea* provides insights into the evolution of secondary plastids. BMC Biology, 16(1). Available from: 10.1186/s12915-018-0593-5

Cock, P.J., Antao, T., Chang, J.T., Chapma, B.A., Cox, C.J., Dalke, A., Friedberg, I., Hamelryck, T., Kauff, F., Wilczynski, B., de Moon, M.J. (2009) Biopython: freely available Python tool for computational molecular biology and bioinformatic. Bioinformatics, 25, 1422–1423. Available from: 10.1093/bioinformatics/btp163

Deane, J.A., Strachan, I.M., Saunders, G.W., Hill, D.R.A. & McFadden, G.I. (2002) Cryptomonad evolution: nuclear 18S rDNA phylogeny versus cell morphology and pigmentation. Journal of Phycology, 38, 1236–1244. Available from: 10.1046/j.1529-8817.2002.01250.x

Douglas, S.E., Penny, S.L. (1999) The plastid genome of the cryptophyte alga, *Guillardia theta*: complete sequence and conserved synteny groups confirm its common ancestry with red algae. Journal of Molecular Evolution, 48, 236–244

Ekelund, F. & Patterson, D.J. (1997) Some heterotrophic flagellates from a cultivated garden soil in Australia. Archiv für Protistenkunde, 148(4), 461–478. Available from: 10.1016/S0003-9365(97)80022-X

Fresenius, G. (1858) Beiträge zur Kenntnis mikroskopischer Organismen. Abhandlungen der Senckenbergischen Naturforschenden Gesellschaft, 211–242. Brönner. Available from: https://books.googleusercontent.com/books/

Guillard, R.L. & Lorenzen, C.J. (1972) Yellow-green algea with Chlorophyllide C1,2. Journal of Phycology, 8, 10–14.

Hall, T.A. (1999) BioEdit: a user-frienldy biological sequence alignment editor and analysis program for Windows 95/98/NT. Nucleic Acids Symposium Series, 41(2), 95–98. Available from: 10.14601/phytopathol_mediterr-14998u1.29

Hernandez-Hernandez, T., Miller, E.C., Roman-Palacios, C. & Wiens, J.J. (2021) Speciation across the tree of life. Biological Reviews, 96, 1205–1242. Available from: doi: 10.1111/brv.12698

Hill, D.R. (1991) Diversity of heterotrophic cryptomonads, In: Patterson, D.J., Larsen, J. (Eds.), The Biology of Free-Living Heterotrophic Flagellates. Clarendon Press, Oxford, England, pp. 235–240.

Hohlfeld, M., Meyer, C., Schoenle, A., Nitsche, F. & Arndt, H. (2022) Biogeography, autecology and phylogeny of percolomonads based on newly described species. Journal of Eukaryotic Microbiology, 70(1), e12930. Available from: 10.1111/jeu.12930

Huelsenbeck, J.P. & Ronquist, F. (2001) MRBAYES: Bayesian inference of phylogenetic trees. Bioinformatics, 17(8), 754–755. Available from: 10.1093/bioinformatics/17.8.754

Jarreta de Castro, A.A. & Bicudo, C.E. (2023) Goniomonas brasiliensis n. sp., In: Flora Ficológica do Estado de São Paulo: Cryptophyceae. Rima Editora, Sao Carlos, pp. 73– 74.

Katoh, K. & Standley, D.M. (2013) MAFFT multiple sequence alignment software Version 7: Improvements in performance and usability. Molecular Biology and Evolution, 30(4), 772–780. Available from: 10.1093/molbev/mst010

Keeling, P.J. (2004) Diversity and evolutionary history of plastids and their hosts. American Journal of Botany, 91(10), 1481–1493. Available from: 10.3732/ajb.91.10.1481

Kim, E. & Archibald, J.M. (2013) Ultrastructure and molecular phylogeny of the cryptomonad *Goniomonas avonlea* n. sp. Protist, 164(2), 160–182. Available from: 10.1016/j.protis.2012.10.002

Kugrens, P. & Lee, R.E. (1991) Organization of cryptomonads, in: Patterson, D.J., Larsen, J. (Eds.), The Biology of Free-Living Heterotrophic Flagellates. Clarendon Press, Oxford, England, pp. 219–233.

Kumar, S., Stecher, G., Li, M., Knyaz, C. & Tamura, K. (2018) MEGA X: Molecular evolutionary genetics analysis across computing platforms. Molecular Biology and Evolution, 35(6), 1547–1549. Available from: 10.1093/molbev/msy096

Larsen, J. & Patterson, D.J. (1990) Some flagellates (Protista) from tropical marine sediments. Journal of Natural History, 24(4), 801–937. Available from: 10.1080/00222939000770571

López-García, P., Rodríguez-Valera, F. & Moreira, D. (2002) Toward the monophyly of Haeckel’s Radiolaria: 18S rRNA environmental data support the sisterhood of Polycystinea and Acantharea. Molecular Biology and Evolution, 19(1), 118–121. Available from: 10.1093/oxfordjournals.molbev.a003976

Martin-Cereceda, M., Roberts, E.C., Wootton, E.C., Bonaccorso, E., Dyal, P., Guinea, A., Rogers, D., Wright, C.J. & Novarino, G. (2010) Morphology, ultrastructure, and small subunit rDNA phylogeny of the marine heterotrophic flagellate *Goniomonas* aff. *amphinema*. Journal of Eukaryotic Microbiology, 57(2), 159–170. Available from: 10.1111/j.1550-7408.2009.00449.x

McFadden, G.I. (2018) Genome of tiny predator with big appetite. BMC Biology, 16(1), 140. Available from: 10.1186/s12915-018-0610-8

McFadden, G.I. (2017) The cryptomonad nucleomorph. Protoplasma, 254(5), 1903–1907. Available from: 10.1007/s00709-017-1153-5

Medlin, L., Elwood, H.J., Stickel, S. & Sogin, M. (1988) The characterization of enzymatically amplified eukaryotic 16S-like rRNA-coding regions. Gene, 71(2), 491–499. Available from: 10.1016/0378-1119(88)90066-2

Mignot, J.-P. (1965) Étude ultrastructurale de (*Cyathomonas truncata*) From. (flagellé Cryptomonadine). Journal de Microscopie, 4, 239–252.

Novarino, G. (2003) A companion to the identification of cryptomonad flagellates (Cryptophyceae = Cryptomonadea). Hydrobiologia, 502(1), 225–270. Available from: 10.1023/B:HYDR.0000004284.12535.25

Patterson, D.J. & Lee, W.J. (2000) Geographic distribution and diversity of free-living heterotrophic flagellates, in: Leadbeater, B.S.C., Green, J.C. (Eds.), In: The Flagellates - Unity, Diversity and Evolution. Taylor & Francis Ltd, London, pp. 269–287.

Phanprasert, Y., Kim, S.Y., Kang, N.S., Jeong, M., Kim, J.I., Shin, W., Lee, W.J. & Kim, E. (2025) Morphological and molecular phylogenetic characterization of three new marine goniomonad species, Journal of Eukaryotic Microbiology, 72:e70002 Available from: 10.1111/jeu.70002

Ronquist, F., Teslenko, M., Van Der Mark, P., Ayres, D.L., Darling, A., Höhna, S., Larget, B., Liu, L., Suchard, M.A. & Huelsenbeck, J.P. (2012) MrBayes 3.2: Efficient bayesian phylogenetic inference and model choice across a large model space. Systematic Biology, 61(3), 539–542. Available from: 10.1093/sysbio/sys029

Schoenle, A., Hohlfeld, M., Rosse, M., Filz, P., Wylezich, C., Nitsche, F. & Arndt, H. (2020) Global comparison of bicosoecid *Cafeteria*-like flagellates from the deep ocean and surface waters, with reorganization of the family Cafeteriaceae. European Journal of Protistology, 73, 125665. Available from: 10.1016/j.ejop.2019.125665

Shiratori, T. & Ishida, K. (2016) A new heterotrophic cryptomonad: *Hemiarme marina* n.g., n. sp. Journal of Eukaryotic Microbiology, 63, 804–812. Available from: doi:10.1111/jeu.12327

Shalchian-Tabrizi, K., Bråte, J., Logares, R., Klaveness, D., Berney, C. & Jakobsen, K. (2008) Diversification in unicellular eukaryotes: cryptomonad colonizations of marine and freshwaters inferred from revised 18S rRNA phylogeny. Environmental Microbiology, 10(10), 2635–2644. Available from 10.1111/j.1462-2920.2008.01685.x

Simek, K., Mukherjee, I., Szoke-Nagy, T., Haber, M., Salcher, M. M., & Ghai, R. (2023) Cryptic and ubiquitous aplastidic cryptophytes are key freshwater flagellated bacterivores. ISME Journal, 17(1), 84–94. Available from: 10.1038/s41396-022-01326-4

Stamatakis, A. (2014) RAxML version 8: a tool for phylogenetic analysis and post-analysis of large phylogenies. Bioinformatics, 30(9), 1312–1313. Available from: 10.1093/bioinformatics/btu033

Stein, F. (1878) Der Organismus der Flagellaten I., In: Der Organismus der Infusionstiere III. Wilhelm Engelmann, Leipzig.

Von Der Heyden, S., Chao, E. & Cavalier-Smith, T. (2004) Genetic diversity of goniomonads: an ancient divergence between marine and freshwater species. European Journal of Phycology, 39(4), 343–350. Available from: 10.1080/09670260400005567

Vørs, N. (1993) Marine heterotrophic amoebae, flagellates and heliozoa from Belize (Central America) and Tenerife (Canary Islands), with descriptions of new species, *Luffisphaera bulbochaete* n. sp., *L. longihastis* n. sp., L. turriformis n. sp. and Paulinella intermedia n. sp. Journal of Eukaryotic Microbiology, 40, 272–287. Available from: 10.1111/j.1550-7408.1993.tb04917.x

Wylezich, C., Meisterfeld, R., Meisterfeld, S. & Schlegel, M. (2002) Phylogenetic analyses of small subunit ribosomal RNA coding regions reveal a monophyletic lineage of euglyphid testate amoebae (Order Euglyphida). Journal of Eukaryotic Microbiology, 49(2), 108–118. Available from: 10.1111/j.1550-7408.2002.tb00352.x

Yabuki, A., Kamikawa, R., Ishikawa, S.A., Kolisko, M., Kim, E., Tanabe, A., Kume, K., Ishida, K.-I., Iangki, Y. (2014) *Palpitomonas bilix* represents a basal cryptist lineage: insight into the character evolution in Cryptista. Scientific Reports, 4, 4641. Available from: 10.1038/srep04641

