## Supplementary tables for "Cryptic Cryptophytes – revision of the genus *Goniomonas*"

[1]University of Cologne. Institute of Zoology. Department of General Ecology and Limnology. Zuelpicher Straße 47b. 50674 Cologne. Germany
[2]Senckenberg am Meer, DZMB – German Centre for Marine Biodiversity Research, Südstrand 44, 26382 Wilhelmshaven, Germany.

Table 1 List of chemicals and their amounts to produce one litre of Schmalz-Pratt Medium with 23 ‰ salinity.

| Chemical component | Amount [g/l] for 23 PSU |
|---|---|
| NaCl | 18.77 |
| KCl | 0.45 |
| $MgCl_2 \times 6\ H_2O$ | 3.67 |
| $MgSO_4 \times 7\ H_2O$ | 4.61 |
| $CaCl_2 \times 2\ H_2O$ | 0.97 |
| $KNO_3$ | 0.1 |
| $K_2HPO_4 \times 3\ H_2O$ | 0.01 |

Table 2 Pairwise distances (in percent) of 18S rDNA sequences for freshwater clusters including the genera *Goniomonas* and *Aquagoniomonas* n. g.

| | Strain | 1 | 2 | 3 | 4 | 5 | 6 | 7 | 8 | 9 |
|---|---|---|---|---|---|---|---|---|---|---|
| 1 | HFCC 235 *Goniomonas truncata* | 0 | | | | | | | | |
| 2 | HFCC 232 *Goniomonas truncata* | 0 | 0 | | | | | | | |
| 3 | HFCC 251 *Goniomonas rhenensis* n. sp. | 3.83 | 3.83 | 0 | | | | | | |
| 4 | HFCC 23 *Aquagoniomonas mylnikovii* n. g. et. n. sp. | 10.47 | 10.47 | 12.65 | 0 | | | | | |
| 5 | LC000677 | 10.49 | 10.49 | 12.55 | 0.47 | 0 | | | | |
| 6 | HFCC 163 "*Aquagoniomonas* sp." | 10.77 | 10.77 | 12.75 | 0.61 | 0.52 | 0 | | | |
| 7 | HFCC 5007 *Aquagoniomonas atacamiensis* n. g. et n. sp. | 10.4 | 10.4 | 12.69 | 3.1 | 0.82 | 0.87 | 0 | | |
| 8 | HFCC 5008 *Aquagoniomonas* sp. | 9.86 | 9.86 | 12.91 | 0.72 | 0.82 | 0.78 | 0.61 | 0 | |
| 9 | AY360459 | 10.71 | 10.71 | 13.11 | 0.98 | 0.51 | 1.03 | 1.42 | 1.32 | 0 |

Table 3 Pairwise distances in percent based on 18S rDNA for the cluster *Limnogoniomonas* n. g.

| | Strain | 1 | 2 | 3 | 4 |
|---|---|---|---|---|---|
| 1 | HFCC 162 *Limnogoniomonas fijiensis* n. g. et n. sp. | 0 | | | |
| 2 | DQ980479 | 0.17 | 0 | | |
| 3 | U03072 | 0.28 | 0.42 | 0 | |
| 4 | HQ659565 | 7.7 | 8.22 | 7.98 | 0 |

Table 4 Pairwise distances (in percent) of 18S sequences for the cluster including the genus *Baltigoniomonas* n. g.

| | Strain | 1 | 2 |
|---|---|---|---|
| 1 | HFCC 22 *Baltigoniomonas juergensii* n. g. et n. sp. | 0 | |
| 2 | MK177625 | 0.31 | 0 |

Table 5 Pairwise distances (in percent) of 18S sequences for the cluster including the genus *Neptunogoniomonas* n. g.

| | Strain | 1 | 2 | 3 |
|---|---|---|---|---|
| 1 | HFCC 1525 | 0 | | |
| 2 | JQ434475 *Neptunogoniomonas avonlea* n. comb. | 0.31 | 0 | |
| 3 | LC647565 | 0 | 0.41 | 0 |

Table 6 Pairwise distances based on 18S rDNA for the cluster *Cosmogoniomonas* n. g.

| | Strain | 1 | 2 | 3 | 4 |
|---|---|---|---|---|---|
| 1 | HFCC 662 *Cosmogoniomonas* sp. | 0 | | | |
| 2 | LC674566 | 0.13 | 0 | | |
| 3 | EU047707 *Cosmogoniomonas martincerecedai* n. comb. | 0.13 | 0.12 | 0 | |
| 4 | AF508277 *Cosmogoniomonas pacifica* n. g. et n. sp. | 3 | 2.57 | 2.57 | 0 |

Table 7 Pairwise distances in percent based on 18S rDNA sequences for the genera *Marigoniomonas* n. g., *Thalassogoniomonas* n. g. and *Poseidogoniomonas* n. g. and the associated sequences.

| | Strain | 1 | 2 | 3 | 4 | 5 | 6 | 7 | 8 |
|---|---|---|---|---|---|---|---|---|---|
| 1 | HFCC 841 *Marigoniomonas* sp. | 0 | | | | | | | |
| 2 | KX431494 | 2.12 | 0 | | | | | | |
| 3 | HFCC 5003 *Marigoniomonas* sp. | 0.26 | 2.21 | 0 | | | | | |
| 4 | HFCC 272 *Marigoniomonas* sp. | 3.4 | 3 | 3.53 | 0 | | | | |
| 5 | HFCC 1666 *Thalassogoniomonas amphinema* n. g. et n. sp. | 6.1 | 4.76 | 6.32 | 6.78 | 0 | | | |
| 6 | AY360454 | 6 | 4.22 | 6.18 | 6.38 | 1.44 | 0 | | |
| 7 | HFCC 157 *Poseidogoniomonas* sp. | 6.09 | 3.87 | 6.27 | 6.53 | 3.73 | 3.33 | 0 | |
| 8 | BEAP0122b | 6.38 | 4.31 | 6.73 | 7.52 | 4.67 | 3.85 | 3.38 | 0 |

Table 8 List of strains isolated and described in the present study. The number of the strains in the Heterotrophic Flagellate Culture Collection Cologne (HFCC), the location and the geographic coordinates of the sampling sites, the habitat and the NCBI accession number of the 18S rDNA sequence are given.

| Species | Isolate (HFCC) | Location of isolation | Latitude/Longitude | Habitat | Accession number (NCBI) |
|---|---|---|---|---|---|
| *Baltigoniomonas juergensii* n. g. et n. sp. | 22 | Kloster, Island Hiddensee, Germany, Baltic Sea | 54°35'37"N/ 13°06'34"E | Brackish water, 8 PSU, muddy sand | XXX |
| *Marigoniomonas* sp. | 841 | Loberia, Atacama, Chile, Southern Pacific (Kelp, Pacific Ocean) | 23°3'13,72"S/ 70°32"58,63"W | Marine, 36 PSU, shore biofilm, | XXX |
| *Marigoniomonas* sp. | 272 | Ribeira Brava, Madeira Island, Portugal, North Atlantic | 32°36`33"N/ 16°53`16"W | Marine, deep sea, 35 PSU, soft sediment (1000m depth) | XXX |
| *Thalassogoniomonas amphinema* n. g. et n. sp. | 1666 | Helgoland Island, Germany, North Sea | 54°11’01’’ N/ 07°53’23’’ E | Marine, 34 PSU, shore biofilm | XXX |
| *Poseidogoniomonas* sp. | 157 | Island of Faial, Azores, Portugal (Atlantic Ocean) | 38°28'25"N/ 28°27'16"W | Marine, 36 PSU, fine sand (450m depth) | XXX |
| *Goniomonas truncata* | 235 | River Rhine, Cologne, Germany | 50°49'34"N/ 6°35'34"E | Freshwater, river snow | XXX |
| *Goniomonas rhenensis* n. sp. | 251 | River Rhine, Cologne, Germany | 50°49'34"N/ 6°35'34"E | Freshwater, river snow | XXX |
| *Aquagoniomonas mylnikovii* n. g. et n. sp. | 23 | Pond in Borok, Jaroslawl, Russia | 58°03'27"N/ 38°15'37"E | Freshwater, pond biofilm | XXX |
| *Aquagoniomonas atacamiensis* n. g. et n. sp. | 5007 | Rio Loa, Atacama, Chile | 21°25'41"S/ 70°03'15"W | Freshwater, river biofilm | XXX |
| *Limnogoniomonas fijiensis* n. g. et n. sp. | 162 | Rainforest pond, Suva, Fiji | 18°08'29"S/ 178°26'29"E | Freshwater, plankton | XXX |
